# Endotome as a Source of Human Peri-Aortic Brown Adipocytes

**DOI:** 10.64898/2026.07.04.735132

**Authors:** Hao Yu, Weiman Xiang, Kexin Teng, Ethel S.K. Ng, Angel Y.F. Kam, Siwakorn Punyawatthananukool, Stephen Dalton, Tianming Wu

## Abstract

Brown adipocytes (BAs) hold therapeutic promise for obesity and metabolic diseases. While interscapular BAs derive from *Pax3*+/*Myf5*+ dermomyotome, peri-aortic BAs are inferred from an unknown *Pax3*+/*Myf5*- somitic origin. Here, we identify human endotome as an *MYF5*-independent source of peri-aortic BAs. Through interrogating public mouse organogenesis and in-house human trunk embryoid single-cell data, we show that the early endotome cells are *MYF5*-independent and are primed by TGF-β-induced epithelial-to-mesenchymal transition. Mechanistically, endotome-to-BA specification requires sequential BMP inhibition and Wnt activation. This roadmap results in UCP1-expressing and metabolically active BAs that transcriptionally resemble *in vivo* peri-aortic BAT. The multipotent endotome cells also give rise to vascular smooth muscle and endothelial cells, offering a self-sufficient source for BAT vasculature. Endotome-derived BAs show accelerated differentiation, reduced heterogeneity, and sustained Wnt activity. Thus, the endotome provides a versatile platform for generating BAs and supporting vasculature, with implications for cell-based therapy and tissue engineering in metabolic disease.

## Introduction

Brown adipose tissue (BAT) uncouples nutrient catabolism from ATP synthesis to dissipate energy as heat, making it a promising therapeutic approach for metabolic disorders.^1–5^ Understanding the developmental origins of BAT is essential for harnessing its regenerative potential, yet the embryonic provenance of brown adipocytes (BA) remains incompletely understood.

Classically, BAT is thought to arise from *Pax3*+/*Myf5*+ dermomyotome precursors, a lineage shared with skeletal muscle.^3,6–10^ While *Myf5*+ precursors contribute to over 80% of interscapular and subscapular BA populations, approximately 60% cervical BAs and, notably, all peri-aortic BAs do not derive from the *Myf5*+ somitic lineage, despite the majority of these cervical and peri-aortic BAs originating from *Pax3*+ somites.^8^ These observations suggests an alternative *Pax3*+/*Myf5*- somitic origin, though the particular somite compartment from which these BAs arise remains unknown.

The developmental origins of aortic perivascular adipose tissue (PVAT) are closely tied to their unique functional properties. Notably, perivascular endothelial and mural cells isolated from stromal-vascular tissue (SVT) have the ability to differentiate into BAs, resembling the peri-aortic depot of BAT, which is reported to be more resistant to ‘whitening’ than interscapular BAT.^11–16^ This perivascular origin prompted us to investigate the recently described endotome, a somite compartment identified by single-cell RNA sequencing (scRNA-seq) of mouse organogenesis and human trunk embryoid models,^17–19^ that serves as a precursor pool for SVT-related vascular endothelium and mural/smooth muscle cells, constituting the aortic vasculature.^20–22^ Mouse lineage tracing using *VE-cadherin* and *Zfp423* drivers has further implicated the endotome in peri-aortic BA genesis.^23–25^ However, the precise identity and developmental trajectory of these BA-committed endotome precursors, and how they diverge from the classical *Myf5*+ dermomyotome lineage, remain elusive.

A key challenge is to define the divergent developmental programs of endotome and dermomyotome that lead to BA genesis, in order to understand their distinct signaling requirements and functional outputs. Here, through analyzing scRNA-seq datasets of mouse somite development alongside our previous human trunk embryoid model (hTEM), we show that *EBF2*+ endotome cells emerge earlier than *MYF5*+ dermomyotome and serve as an *MYF5*-independent source of BAs. Mechanistically, we identify TGF-β-induced epithelial-to-mesenchymal transition (EMT) as a necessary step for endotome specification toward BA fate, and reveal a distinct signaling logic: whereas dermomyotome-derived BAs require BMP activation and Wnt inhibition,^10^ endotome-derived BA genesis requires sequential BMP inhibition followed by Wnt signaling activation. Comparative transcriptomic integration indicates that *in vitro*-derived endotome cells represent the *in vivo* counterpart of the *PAX3*+/*MYF5*- somitic origin of peri-aortic BAT. We further validate the multilineage potency of endotome precursors toward vascular endothelial (VE) and smooth muscle cells (vSMC). Collectively, our findings provide an efficient strategy to generate the essential components for reconstituting an *in vitro* BA vasculature, with broad implications for metabolic regenerative medicine.

## Results

### Endotome emerges before dermomyotome and correlates with TGF-β-mediated epithelial-to-mesenchymal transition

To understand endotome as a *MYF5*-independent source of BA precursors, we first need to characterize its developmental timing and molecular identity. During mouse somite compartmentalization, the ventral somite undergoes an secondary epithelial-to-mesenchymal transition (EMT) in response to signals from surrounding tissues, generating a ventral mesenchymal layer (source of ventrolateral endotome and ventromedial sclerotome) and a dorsal epithelial dermomyotome.^20,26,27^ By re-analyzing the mouse organogenesis atlas (E7.5–E8.5) from Imaz-Rosshandler et al. study, we identified distinct clusters corresponding to pan-somite (*Pax3, Meox1*), endotome (*Ebf2, Tagln, Kdr, Bmp5*), dermomyotome (*Myf5/6, Pax7*), and sclerotome (*Pax1*) (Figures S1A-S1C).^17–19,21^ After splitting the somitic cells by embryonic days, we found that *Ebf2* labeled the earliest emergence of endotome at E7.75, whereas *Myf5*-expressing mature dermomyotome cells appeared later at E8.5 (Figures 1A-1C). A similar endotome-like compartment (*Ebf2*, *Sim1*) has be found in chick embryos prior to dermomyotome formation.^27^ Notably, cells expressing *Tagln* at E8.5 develop into thermogenic adipocytes,^28^ and Ebf2 is also a master regulator of BA genesis of somite origin.^29,30^ These data suggest that the endotome forms earlier than dermomyotome, represents a distinct somite compartment, and may serve as a precursor pool for BA genesis.

**Figure 1.**
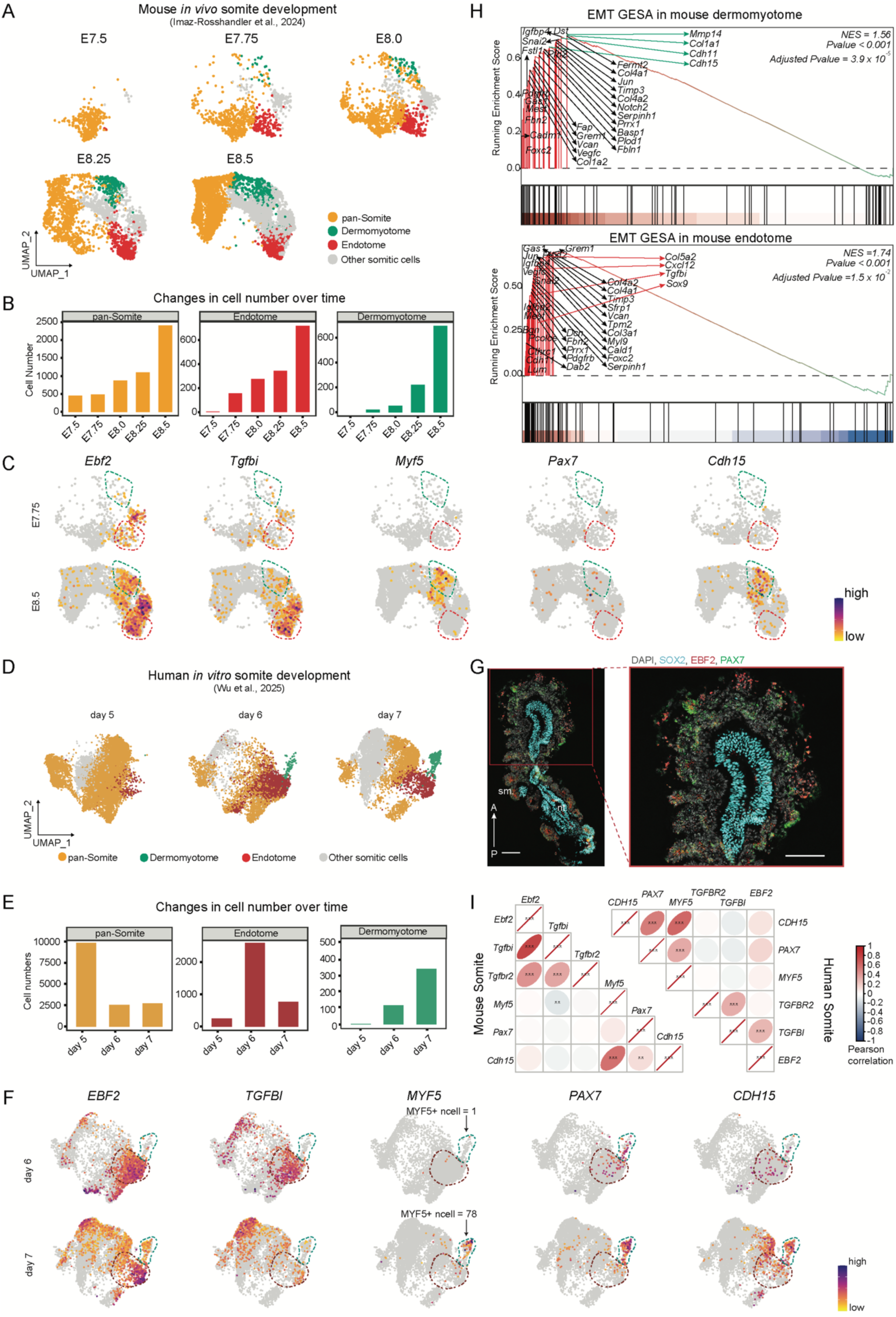
Endotome emerges before dermomyotome and correlates with TGF-β-mediated epithelial-to-mesenchymal transition. (A) UMAP trajectory of somite compartmentalization in mouse E7.5-E8.5 dataset from Imaz-Rosshandler et al., 2024. (B) Changes in cell number over time, corresponding to (A). (C) Featureplot showing the expression levels of indicated genes in E7.75 and E.85 somites. Green dashed line, cluster of dermomyotome. Red dashed line, cluster of endotome. (D) UMAP trajectory of somite compartmentalization in human trunk embryoids (hTEM.v3) (days 5-7 subset) dataset from Wu et al., 2025. (E) Changes in cell number over time, corresponding to (D). (F) Featureplot showing the expression levels of indicated genes in somites from hTEM.v3 at day 6 and 7. Green dashed line, cluster of dermomyotome. Red dashed line, cluster of endotome. (G) Immunohistochemical staining with antibodies against EBF2 (endotome), PAX7 (dermomyotome) and SOX2 (neural tube) in longitudinal sections of day-6.5-hTEM.v3. sm., somite. nt., neural tube. n > 3. Scale bars, 100 µm. (H) Gene set enrichment analysis (GSEA) for differentially expressed gene sets from mouse dermomyotome (top) and endotome (bottom). NES, normalized enrichment score. (I) Heatmap depicting the expression correlation matrix among selected endotome (*EBF2*, *TGFBI*, *TGFBR2*) and dermomyotome (*MYF5*, *PAX7*, *CDH15*) markers. Transcripts in cells were extracted from mouse and human somites in Figures S1A and S1E, respectively. Color represents Pearson correlation. **p < 0.01, ***p < 0.001.

We next turned our focus to the human counterpart of endotome, as somite-derived endotome cells were recently reported in human pluripotent stem cell-derived trunk embryo models.^17,18,31^ Using our previous hTEM version 3 (hTEM.v3), we recapitulated early somite compartmentalization events at days 5-7, corresponding to mouse E7.75-E8.5 (Figure S1E-S1G).^18^ Consistent with the mouse data, human *in vitro* endotome arose at day 6, preceding *MYF5*-expressing dermomyotome at day 7 (Figures 1D-1F). Immunostaining of day-6.5 hTEM.v3 sections confirmed the reciprocal developmental trajectories of endotome and dermomyotome, marked by mutually exclusive expression of EBF2 and PAX7 at protein levels, along the anterior-posterior axis (Figure 1G). Extended studies in mice have established the roles of Wnt and BMP pathways in dermomyotome and subsequent skeletal muscle specification.^32–35^ Consistently, gene ontology (GO) analysis confirmed that top 100 differentially expressed genes in dermomyotome cells were enriched for canonical Wnt and BMP pathways (Figure S1D). In contrast, genes enriched in mouse endotome are associated with BMP and TGF-β pathways (Figure S1D), suggesting a compartment-specific signaling requirement.

Somites undergo secondary EMT processes for distinct compartment formations, controlled by various signaling pathways including Wnt, BMP, TGF- β, PDGF, VEGF and HGF.^36^ We therefore performed gene set enrichment analysis (GSEA) of EMT pathway genes in mouse dermomyotome and endotome cells (Figure 1H). Among the top-ranked EMT genes enriched in dermomyotome, we observed several EMT regulators involved in dorsal somite patterning and skeletal muscle specification (*Col1a1*, *Cdh11*, *Cdh15*, *Mmp14*).^37–40^ Among these, *Cdh15* (encoding M-Cadherin) is a well-known EMT regulator mediating dermomyotome patterning and a target of Wnt signaling.^37,38^ Despite the shared EMT regulators (*Snai2*, *Igfbp4*, *Timp3*), mouse endotome displayed a unique EMT profile (*Col5a2*, *Cxcl12*, *Sox9*, *Tgfbi*), consistent with its ventrolateral location (*Col5a2*, *Sox9, Sim1*) and its involvement in vascular endothelial specification (*Cxcl12*).^21,41^ Importantly, *Tgfbi* (encoding a TGF-β-induced secreted extracellular matrix protein) is significantly enriched in mouse endotome. This aligns with the enrichment TGF-β signaling pathway genes in endotome cells (Figures 1H and S1D).

Finally, hierarchical clustering analysis of mouse and human somite cells revealed distinct correlations between *MYF5*/*PAX7* and *CDH15* (target of Wnt signaling), and between *EBF2* and *TGFBR2*/*TGFBI* (receptor and target *of* TGF- β signaling), respectively (Figure 1I), pointing to divergent secondary EMT processes in dermomyotome and endotome cells. BA progenitors undergo lineage commitment by the time Ebf2 protein is detected in various BAT depots in mice.^30,42,43^ The high expression level of *EBF2* suggests the possibility of endotome as a source of BA genesis. In summary, we delineated the somite developmental sequences of the early endotome and late dermomyotome, pointing to a mechanistic link between TGF-β-induced EMT and endotome patterning that implicates the endotome as BA lineage precursors.

### Directed differentiation of endotome cells with BA-committing competency

Much less is known about the BA genesis potential of endotome precursors, largely due to the lack of suitable cell models. We therefore sought to generate EBF2+ endotome cells from somitic origin. In aim of this, we first needed to achieve high-efficient PAX3+ somite differentiation from human embryonic stem cells (hESC). We previously observed that, in human gastruloids and our previous hTEMs,^44,45^ the starting size of hESC spheroids affects the expression scaling in somite and neural fated cells. In light of this, we treated size-controlled hESC spheroids with the Wnt agnost CHIR-99021 (CHIR), the BMP inhibitor LDN-193189 (LDN) and pro-proliferation supplements including insulin-transferrin-selenium (ITS) and fetal bovine serum (FBS) for 6 days (Figure 2A).^3,10,46^ To monitor cell differentiation toward the PAX3+ somite fate, we used a PAX3-mClover3/EBF2-mScarlet dual reporter H9-hESC line.^18^ By day 6, 80.9 ± 3.9 % cells expressed somite marker PAX3-mClover3 from smaller (100-150 µm diameter) hESC spheroids (Figures 2B and S2A-S2C). Conversely, larger (200-250 µm) hESC spheroids supported both somite (*PAX3*, *SIX1*) and neural (*SOX2*, *PAX6*) fate patterning (Figures S2B). In addition, EBF2 and TGFBI (mesenchymal marker) proteins were observed at the outer cell layer of somites, in contrast to the inner domain of epithelialized somitomeres expressing PAX3 and ZO-1 (epithelial marker) (Figure 2B). Remarkably, after withdrawing CHIR, EBF2-mScarlet was robustly (87.8% ± 5%) expressed at day 8 from smaller PAX3+ somites (Figures 2C and S2D). We also noted that replacing LDN with 10 ng/mL BMP4 at days 6-8 significantly reduced EBF2 expression by ∼40% (Figures S2E and S2F). This agrees with the critical role of BMP inhibition in the ventrolateral somite compartment patterning and the ventrolateral localization of endotome.^27,41,47^

**Figure 2.**
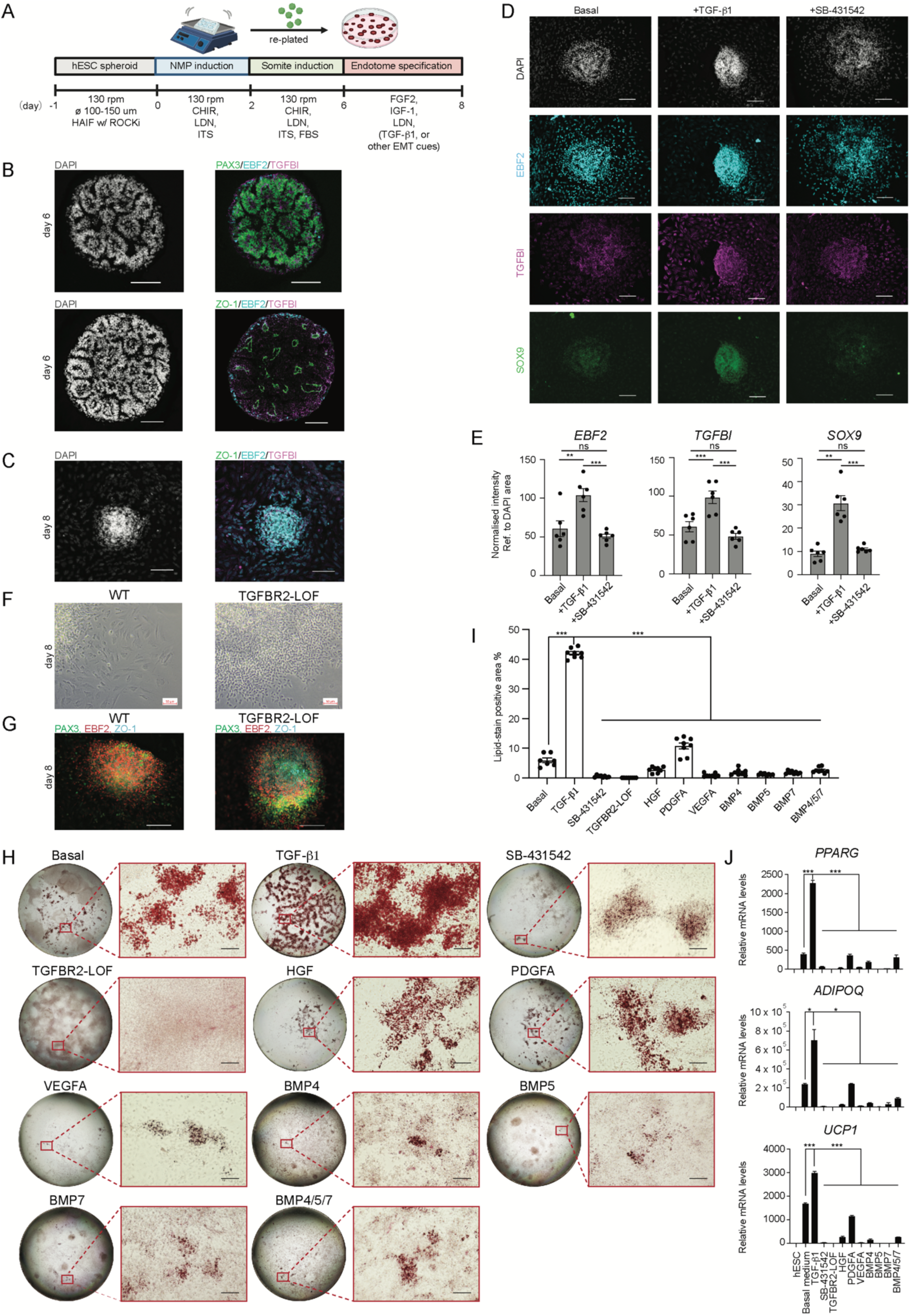
TGF-β-induced EMT is required for BA commitment in endotome cells. (A) Schematic illustrating the generation of endotome cells from 3D hESC cultures. (B) Immunohistochemical staining of hESC-derived somite spheroids with antibodies against PAX3 (somite marker), EBF2 (endotome marker), ZO-1 (epithelial marker) and TGFBI (mesenchymal marker). Scale bars, 100 µm. n > 3. (C) Immunostaining results showing the epithelial (ZO-1)-to-mesenchymal (TGFBI) transition in day-8 endotome cells (without TGF-β1). Scale bars, 100 µm. n > 3. (D) Immunostaining with antibodies against SOX9, EBF2 and TGFBI on day-8 endotome cells under three conditions: untreated (basal), treatment with 10 ng/ml TGF-β1, or exposed to 10 µM SB-431542. n = 6 for each condition. (E) Quantification of fluorescence intensities among conditions in (D). Data were presented as mean ± SEM. ns, no statistical significance. **p < 0.01, ***p < 0.001. (F) Bright-field images of day-8 endotome cell (treated with 10 ng/ml TGF-β1) morphology using wild-type (WT) and TGFBR2-LOF H9-hESC lines. Scale bars in top panel, 50 µm. Scale bars in bottom panel, 100 µm. (G) Immunostaining for PAX3, EBF2 and ZO-1 in day-8 endotome cultures (treated with 10 ng/ml TGF-β1) using wild-type (WT) and TGFBR2-LOF H9-hESC lines. Scale bars, 100 µm. (H) Lipid (Oil Red O) staining results showing lipid formation on day 29 in endotome cells under various EMT induction (TGF-β1, HGF, PDGFA, VEGFA, BMP4/5/7) or inhibition (SB-431542, TGFBR2-LOF) conditions. Zoomed views exhibited lipid droplets formation in respective condition. Experiments were reproduced in more than three biological batches. Scale bars, 100 µm. (I) Quantification of lipid-stain positive areas in BA cultures in (H). n = 8. Data were presented as mean ± SEM. ***p < 0.001. (J) Real-time qPCR analysis of *PPARG*, *ADIPOQ* and *UCP1* in BA cultures as shown (H). Data were presented as mean ± SEM. n = 3. *p < 0.05, ***p < 0.001.

To determine whether EBF2-expressing endotome precursors can directly give rise to UCP1-expressing BAs, we created an EBF2-mScarlet/UCP1-tagGFP H9-hESC line to track adipogenic progression (see Method section). Exposing day-8 endotome cells to a minimal BA induction medium containing triiodothyronine, IBMX, TGF-β inhibitor SB431542 (SB), dexamethasone, FGF2, IGF-1, and rosiglitazone yielded a clear EBF2-to-UCP1 shift between days 30-34 (Figure S2G). In BA cultures examined at day 34, sorted EBF2-mScarlet+ (35 µm mesh pre-filtered to remove lipid-rich BAs) expressed BA progenitor markers including *PPARG*, *CEBPD* and *UCP1* (Figure S2H).^48,49^ Encouragingly, these data confirmed lineage competence of endotome cells toward a BA identity. However, the overall BA differentiation efficiency remained suboptimal. We reasoned that critical developmental cue(s) was missing for fully committing endotome cells to the BA fate.

### TGF-β-induced EMT primes endotome cells for BA genesis

It was previously believed that TGF-β signaling negatively affects BA progenitor specification at late stages, from E12.5 to E15.5 in mice.^3,10,50–52^ However, those studies focused primarily on the interscapular BAT depot and effect of TGF-β signaling on BA genesis at a fibroblastic stage between E12.5-E15.5, whereas the TGF-β signaling in endotome-to-BA specification was implicated at earlier stages (E7.75-E8.5 in mice and Carnegie stages 9-10 in humans) (Figure 1). To address the role of TGF-β signaling in endotome-derived BA genseis, we first assessed the responsiveness of endotome cells to 10 ng/mL TGF-β1 (ligand) or 10 µM SB (receptor inhibitor) between days 6-8 (Figure 2A). Upon treatment with TGF-β1, protein levels of EBF2, TGFBI and SOX9 (mesenchymal marker in ventral somite) were significantly upregulated, and this upregulation can be suppressed by SB (Figures 2D and 2E). Given the positive correlation between *TGFBR2* (TGF-β receptor) and both *EBF2* and *TGFBI* (TGF-β target) in endotome (Figure 1H), we generated a *TGFBR2* loss-of-function (TGFBR2-LOF) H9-hESC line to further validate the mechanistic role of TGF-β signaling in endotome patterning and subsequent BA commitment (see Method section). Using this TGFBR2-LOF line, we generated day-8 endotome cells in presence of TGF-β1. Endotome cells of TGFBR2-LOF retained the epithelial somite cell markers PAX3 and ZO-1 and exhibited a spheroidal shape, in contrast to wild-type endotome cells, which displayed a mesenchymal-like spindle shape (Figures 2F and 2G). These results indicate that positive feedback of TGF-β signaling mediates secondary EMT during somite-to-endotome differentiation (Figure 1 and Figures 2A-2D).^47,53,54^

To determine whether TGF-β activation is required for endotome cell commitment to BA fate, we reassessed their brown adipogenic potential after TGF-β1 pretreatment and compared it with other signaling inducers involved in somite secondary EMT processes, including PDGFA, VEGFA, HGF and BMP4/5/7 (Figure 2H).^36^ Strikingly, among all EMT inducers tested, TGF-β1 treatment yielded the highest (42 ± 1.7%) BA induction efficiency, as quantified by Oil Red O staining and real-time quantitative PCR of *UCP1*, *ADIPOQ*, and *PPARG* in day-29 cultures (Figures 2I and 2J). Conversely, inhibition of TGF-β pathway with SB or genetic ablation of *TGFBR2* using TGFBR2-LOF line completely abrogated BA differentiation. BMP4 and BMP7 are well-known brown adipogenic ligands abundantly expressed by late-stage fibroblastic pre-BA progenitors in the interscapular BAT depot.^10,55–57^ However, strikingly, substituting the BMP inhibitor LDN with BMP ligands (BMP4/5/7 individually or in combination) at days 6-8 fully abolished lipid production and *UCP1* expression by day 29 (Figures 2F and 2G). These findings suggests that endotome-derived BA genesis is governed by a regulatory logic distinct from that in the interscapular BA lineage.^3,10,55,57^ We further reasoned that the endotome-specific expression of *BMP5* likely acts in a paracrine manner to influence neighboring cells, rather than cell-autonomously affecting its own BA differentiation potential (Figures S2C and S2G).^35^ Collectively, these data established that TGF-β activation and BMP inhibition are required for maintaining the *EBF2*-expressing endotome identity and driving its progression towards a BA fate (Figures S2E, S2F, 2H and 2I).

### Characterization of endotome cells as precursors of BA progenitors

To further characterize the differentiation of endotome cells toward BA fate, we performed scRNA-seq on day-6 somite and day-8 endotome cultures (all endotome cells were treated with TGF-β1 hereafter) using the Singleron platform (Figure 2A).^58^ 13,678 and 11,228 cells from day-6 and day-8 samples were profiled, respectively (Figure 3A). Consistent with the PAX3-mClover3+ flow cytometry results (Figures S2A-S2C), 92.7% of day-6 cells belong to a large somite cluster expressing *PAX3*, *MEOX1* and *SIX1* (Figures 3B-3D). Clustering analysis of day-8 scRNA-seq data resolved detailed subtypes within the major *EBF2*-exprssing pan-endotome population at day 8, including endotome progenitors (End. Pro., 12.7%, *PAX3^high^*/*EBF2^high^*), endotome (29.4%, *PAX3^low^/EBF2^high^*, *SOX9*, *ST8SIA6*, *PLVAP*, *LBH*), endotome-vascular smooth muscle progenitors (End.-vSMC, 40.4%, *MYH11*, *MYO9B*, *PENK*, *ERG*) and endotome-vascular endothelial progenitors (End.-VE, 17%, *TAGLN, TGFBR1, NTM*, *CDH5*) (Figures 3A, 3C and 3D).^43,59^ A small cluster of VE (*SOX17*, *CD34, KDR*) was also observed (Figure 3D). In human interscapular BAT and PVAT, adipogenic fibroblastic precursors and mesenchymal stem cells (MSC) are signified by surface markers such as CD73 (*NT5E*), CD90 (*THY1*), CD105 (*ENG*) and CD146 (*MCAM*) (Figures S3A and S3B).^10,13,56,60,61^ However, *CD73* mRNA and CD73/CD105 proteins were undetectable in day-8 endotome cultures (Figures S3C and S3D). Moreover, the terminal BA progenitor marker *PPARG* was also not absent (Figure 3D).^30,48^ Instead, the abundant expression of vascular-associated genes such as *TAGLN*, *PENK*, *PLVAP* and *ERG*, suggested that the early differentiation potential of endotome cells extends beyond the BA lineage. This aligned with the proposed contribution of endotome cells to aortic VE and vSMC populations in the trunk region,^19,20,47^ where *Tagln*, *Penk*, *Plvap* and *Erg* have been used to track specific vSMC and VE subtypes in peri-aortic vasculature.^43,62,63^ Further lineage tracing studies will determine the early contribution of endotome cells to BA vasculature.

**Figure 3.**
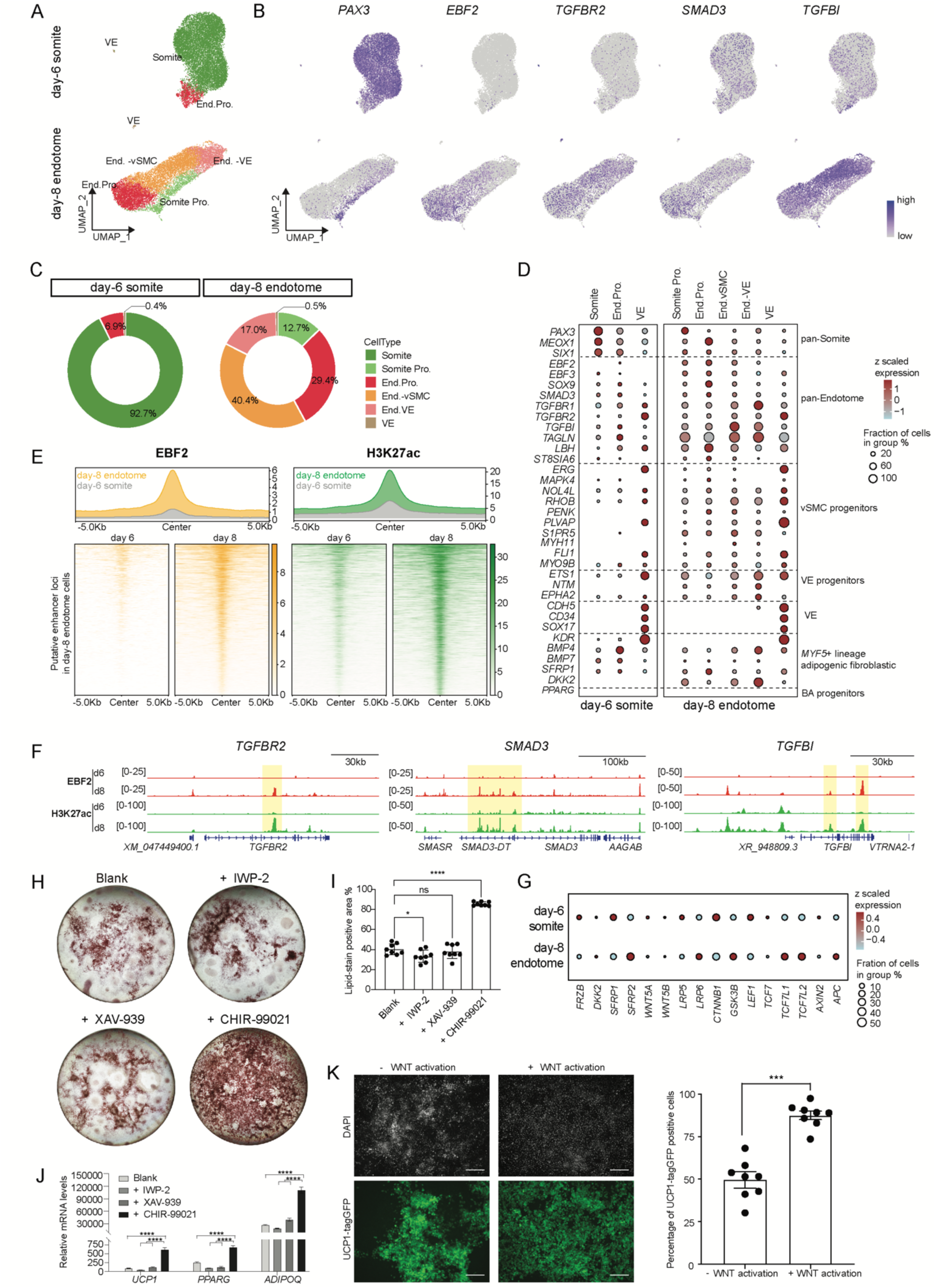
Wnt activation completes BA fate commitment of endotome cells. (A) UMAP showing the identified subtypes of (top) day-6 somite and (bottom) day-8 endotome cells. End. Pro., endotome progenitors. VE, vascular endothelial cells. vSMC, vascular smooth muscle cell. (B) Featureplots showing representative markers of somite identity (*PAX3*), endotome identity (*EBF2*) and TGF-β signaling pathway (*TGFBR2*, *SMAD3*, *TGFBI*). (C) Stack plots showing the proportions of identified cell types in (left) day-6 somite and (right) day-8 endotome cultures. Colors and annotations are the same as in (A). (D) Dotplot displaying a curated list of marker genes delineating identified cell types in (A). Fraction of detected cells below 1% were omitted. (E) Mean (top) and rank-ordered (bottom) CUT&Tag signal intensities for EBF2 and H3K27ac at putative enhancer regions in endotome cells. (F) Genome browser tracks showing CUT&Tag signals for EBF2 and H3K27ac (putative enhancer marker) in day-6 somite and day-8 endotome cells. Shaded areas in yellow highlighted EBF2-bound putative enhancers in endotome cells. (G) Dotplot showing scaled expression of core WNT pathway genes in day-6 somite (d6) and day-8 endotome (d8) cells. Cells analyzed were from (A). (H) Lipid (Oil Red O) staining results showing lipid formation on day 33, under conditions where endotome cells were subjected to 4 days of WNT activation (CHIR-99021) or inhibition (IWP-2 and XAV-939). Blank control was treated with basic medium containing FGF2 and ITS only. Experiments were reproduced in more than three biological batches. (I) Quantification of lipid-stain positive areas in BA cultures in (H). n = 8. Data were presented as mean ± SEM. *p < 0.05, ***p < 0.001. ns, no statistical significance. (J) Real-time qPCR analysis of *UCP1*, *PPARG* and *ADIPOQ* in BA cultures as shown (H). Data were presented as mean ± SEM. n = 3. *p < 0.05, ****p < 0.0001. (K) Left, representative images of UCP1-tagGFP in day-33 BA cultures with (CHIR-99021) or without (blank) 4 days of WNT activation. Right, fraction of UCP1-tagGFP positive cells in relative to DAPI stained cells. n = 8. Data were presented as mean ± SEM. ***p < 0.001.

We next explored endotome-specific transcriptional program underlying the EMT process. Most pan-endotome cells upregulated mRNAs of *TGFBR2* (TGF-β receptor), *TGFBI* (TGF-β target) and *EBF2*, while downregulated *PAX3* (Figure 3B). These dynamics are consistent with TGF-β-induced secondary EMT process we observed in Figures 1 and 2. Furthermore, day-8 endotome cells were enriched for various EMT-associated genes that regulate cytoskeleton and cell motility, including *FOXS1 (*TGF-β target), *S1PR5* (sphingosine-1-phosphate receptor 5), *RHOB* (Ras homolog family member B), *MAPK4* (mitogen-activated protein kinase 4) and *NOL4L* (nucleolar protein 4-like) (Figure 3D).^64–68^ To gain more mechanistic insights, we performed CUT&Tag using EBF2 and H3K27ac (putative enhancer marker) antibodies on cells of day-6 and day-8 cultures. After intersecting H3K27ac-associated genes with the top1000 differentially expressed genes generated from scRNA-seq dataset, we identified 4,052 putative enhancers that were unique to endotome cells, where both EBF2 and H3K27ac signals were substantially increased at day 8 (Figure 3E). Genome browser tracks showed EBF2 occupancy at putative enhancers around representative target genes in endotome cells, including (1) components of TGF-β signaling pathway (*TGFBR2*, *SMAD3*, *TGFBI, FOXS1*); (2) EMT regulators (*S1PR5*, *FOXS1*, *NOL4L*, *PLVAP*, *RHOB*); and (3) endotome-enriched genes driving vSMC and VE specifications (*TAGLN*, *FLI1*, *PDGFRA*) (Figures 3F and S3E). In contrast, EBF2 and H3K27ac signals were absent at loci associated somite induction (*PAX3*), late-stage adipogenic fibroblasts (*BMP7*) and terminal BA progenitors (*PPARG*) in day-8 endotome cells (Figure S3E).^10^

Interestingly, although GATA6 is a classical TGF-β target and has recently been reported as a crucial regulator of BA genesis in the dermomyotome-derived BA lineage,^10,51^ we did not detect GATA6 protein in day-8 endotome cells co-stained with antibodies against EBF2 and SMAD2/3 (Figure S3F). Consistently, EBF2-bound enhancers were absent at *GATA6* locus (Figure S3E). Thus, endotome and dermomyotome lineages differ in their dependency on GATA6 during BA fate commitment.

Collectively, these transcriptomic and epigenetic data, combining with our functional observations (Figure 2), we establish a reciprocal regulatory circuit in which TGF-β signaling and EBF2 cooperatively orchestrate the EMT program to prime endotome cells for BA specification. Nevertheless, the absence of *PPARG* expression at this stage (Figure 3D) and suboptimal (∼40%) BA induction efficiency (Figures 2H) suggests that additional signaling pathway and developmental process are required to complete BA commitment.

### Wnt activation completes BA fate commitment of endotome cells

During somite compartmentalization, Wnt signaling is important for driving dorsal somite (e.g., dermomyotome) lineage, while repressing ventral somite (e.g., sclerotome) fates.^69^ However, the role of Wnt signaling in the ventrolateral endotome compartment is still unknown. In mouse somite development and our hTEM.v3, *FRZB* (encoding a Wnt antagonist) was highly expressed in pan-somite cells, but absent in subsequently formed endotome cells (Figure S3G),^18,19^ implying a de-repression of Wnt signaling during endotome patterning. Consistently, *FRZB* was downregulated in day-8 endotome cells, accompanied with expression of Wnt targets (*DKK1*/*2*, *SFRP1*/*2*), receptors (*FZD10*, *LRP5*/*6*) and core pathway elements (*CTNNB1*, *GSK3B*, *TCF7L1*/*2*, *APC*) (Figure 3G). Wnt inhibition was previously reported to be required for interscapular BA development in mice.^70^ However, the observation of Wnt signaling pathway gene expression in endotome culture raised the possibility that Wnt activation may be involved in endotome-derived BA development.

To test this idea, following BMP inhibition and TGF-β activation, we further allowed Wnt signaling activation by adding 3 µM CHIR for 4 days prior to BA induction. Remarkably, the resultant day-33 BA culture displayed a 2-fold (85.7 ± 2 %) increase in the lipid-filled BA population, compared to the blank control (40.3 ± 5.5 %) (Figures 3H and 3I). Importantly, *PPARG* mRNA was strongly induced after Wnt signaling activation (Figure 3J). In agreement with the lipid staining data, analysis of the UCP1-tagGFP line showed that Wnt activation resulted in 87.5 ± 2.5 % UCP1-exprssing BA cells on day 33, compared with 49.6 ± 4.1% in the blank control (Figure 3K). In contrast, Wnt inhibition by 10 µM IWP-2 (Porcupine inhibitor) or 10 µM XAV-939 (Trankyrase inhibitor) abolished the BA genesis (Figures 3H and 3I), evident by the downregulation of *UCP1*, *PPARG* and *ADIPOQ* at mRNA levels (Figure 3J). Altogether, these results demonstrate that TGF-β activation and BMP inhibition followed by Wnt activation fully potentiate endotome cells to develop into BAs. This is in contrast to the requirement for BMP activation and Wnt inhibition in committing dermomyotome lineage toward BA fate.^10,55,70^

### Comparative analysis of *in vitro* BA development from endotome and dermomyotome lineages

Having established the temporal and signaling relationship between the endotome and dermomyotome BA lineages, we next sought to determine whether the endotome gives rise to the *PAX3*+/*MYF5*- originated peri-aortic BAs. To this end, we performed stepwise, lineage-divergent differentiation of BAs from the two somite compartments *in vitro*. As a control, we concurrently generated dermomyotome-derived interscapular BAs leveraging a modified protocol from previous studies (Figure 4A).^3,10,71^ Briefly, following somite induction on day 6, a 2-day treatment with 3µM CHIR, 10 ng/ml HGF and 2 ng/ml IGF-1 led to the differentiation of *MYF5*/*PAX7*+ dermomyotome at day 8 (Figures 4A and S4A). Subsequent scRNA-seq of day-8 dermomyotome cells and integration with day-6 somite and day-8 endotome datasets confirmed divergent somite compartmentalization trajectories, the dermomyotome precursors displayed developmental potential toward skeletal muscles (SKM); whereas the endotome precursors exhibited potential toward vSMCs (Figures S4B-S4G). Consistent with the developmental sequence of interscapular BAs,^10^ continued culture with HGF and IGF-1 promoted the differentiation of PAX7+ precursors by day 12, followed with a gradual increase in SKM markers (*MYF5*, *MYOD1*, *MYOG*, *MYH3*/*7*) at days 12-22 (Figures S4H and S4I).^71^ By day 26, immunofluorescence results validated the emergence of GATA6+ BA precursors alongside SKM maturation, the latter of which was confirmed by the ubiquitous immunostaining of PAX7, myogenenin (MYOG), myosin heavy chains and Titin (Figures S4J and S4K).^10,51^ BAs of dermomyotome was subsequently differentiated using standard BA induction medium at days 26-46 (Figure 4A). Notably, mRNA and protein of both PAX7 and GATA6 were not expressed in the endotome-BA lineage, at the corresponding BA precursor days 12 and 26 (Figures S4H and S4J), highlighting a fundamental difference between endotome- and dermomyotome-derived BA genesis.

**Figure 4.**
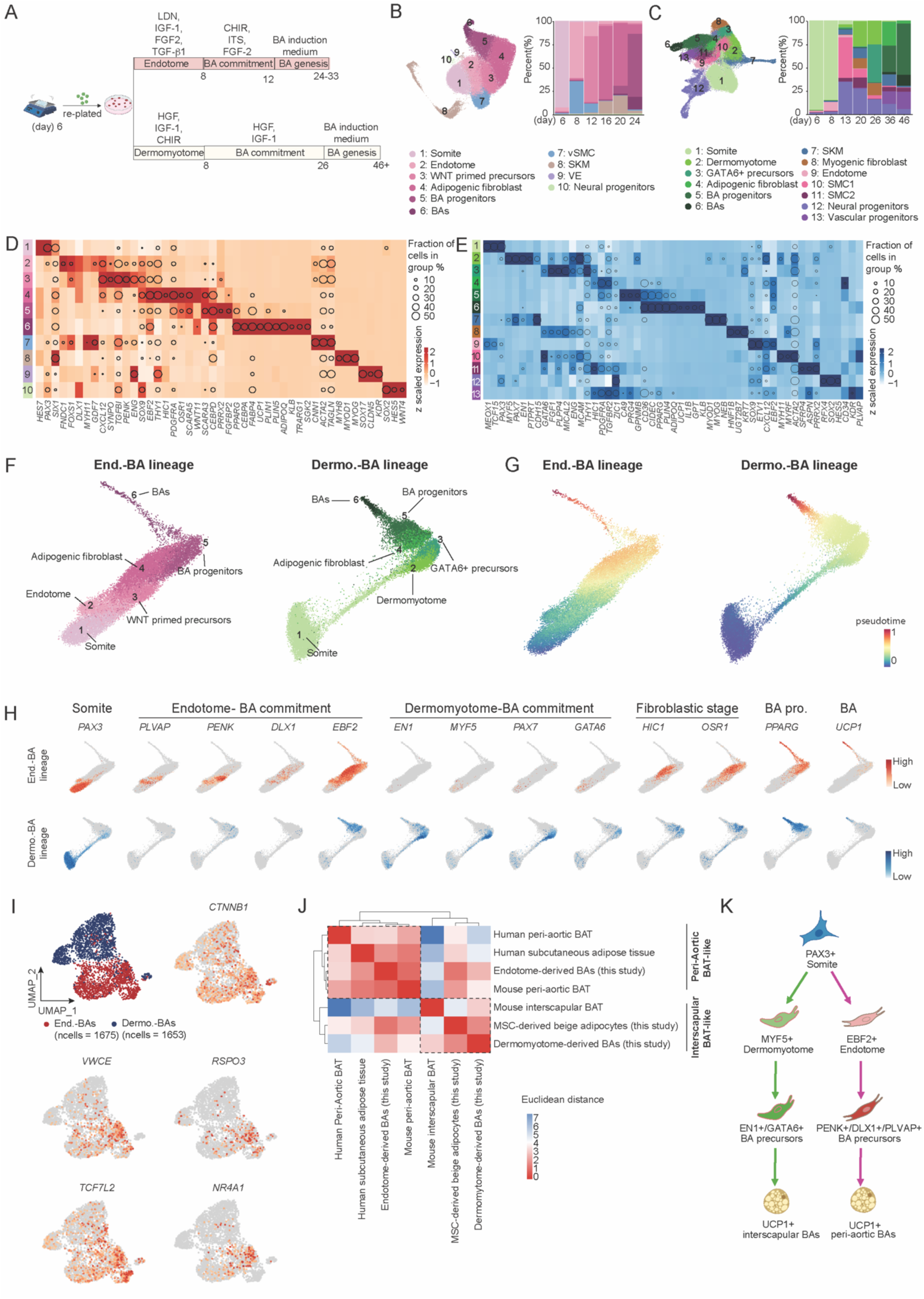
Comparative analysis of *in vitro* BA development from endotome and dermomyotome lineages. (A) Schematic depicting the parallel BA differentiation from somite-derived endotome and dermomyotome cultures. (B) Left, UMAP embedding of identified cell types across BA development of endotome (day 6, 8, 12, 16, 20, 24). Right, corresponding stack plot showing fractions of cellular components across time. (C) UMAP embedding of identified cell types across BA development of dermomyotome (day 6, 8, 13, 20, 26, 36, 46). Right, corresponding stack plot showing fractions of cellular components across time. (D) Dotplot showing scaled expression of a curated list of marker genes in identified cell types in (B). Fraction of detected cells below 1% were omitted. (E) Dotplot showing scaled expression of a curated list of marker genes in identified cell types in (C). Fraction of detected cells below 1% were omitted. (F) Force directed graph visulization of lineage-resolved cells engaged in BA genesis, with endotome-derived (left) and dermomyotome-derived (right) populations shown separately. (G) Force directed graph showing extracted cell embeddings with of pseudotime scores, inferring the developmental progression of cells shown in (F). (H) Featureplots showing the expression of lineage-specific genes marking staged BA development in endotome (top) and dermomyotome (bottom) cells from (F). (I) Featureplots showing the divergent expression pattern of *VWCE* (WNT activator), *RSPO3* (WNT activator), *TCF7L2* (WNT effector) and *NR4A1* (WNT target) in BAs derived from endotome and dermomyotome cultures. (J) Hierachical clustering heatmap comparing BAs derived from endotome and dermomyotome cultures to other BA datasets. Color scale (0 - 7) represents Euclidean distance between samples. Public datasets for this analysis include human subcutaneous adipose tissues (GSE176171), human and mouse peri-aortic BAT (GSE164528), and mouse interscapular BAT (GSE185623). (K) Schematic summary of distinct peri-aortic and interscapular BA genesis stages in endotome and dermomyotome lineages, respectively.

To further characterize the developmental trajectories leading to BAs, we performed time-course scRNA-seq and integrated cells from divergent BA genesis cultures through endotome (total of 72,927 cells, at days 6, 8, 12, 16, 20, 24) and dermomyotome (total of 81,800 cells, at days 6, 8,13, 20, 26, 36, 46) lineages, respectively (Figures S5A-S5D). Louvain-based clustering resolved major somite-derived cell populations that recapitulate the full BA differentiation program of both lineages (Figures 4B-4E). Compared with our dermomyotome-BA cultures as reported by Rao et al., 2023,^10^ endotome-BA cultures exhibited less heterogeneity and fewer non-target cell populations, such as neural and muscle cells (Figures 4B and 4C). A declining fraction of *PAX7*+/*MYOG*+ SKM-fated progenitors was observed in endotome-BA cultures between days 16-24 (Figures 4B, 4D and S5B). These *PAX7*+/*MYOG*+ cells likely originated from spontaneous dermomyotome induction resulting from Wnt activation in residual PAX3+ somite progenitors present at days 8-12.^33^ Importantly, we ruled out their contribution to the endotome-BA trajectory (clusters 2-6), as the interscapular BA lineage tracer *EN1* was exclusively expressed in dermomyotome-BA trajectory (clusters 2-6) (Figures 4E, 5E and 5F).^10,34^

To dissect the molecular pathways underlying the divergent developmental processes toward BAs, we extracted parallel cell clusters 1-6, which represents end-to-end trajectories from the two somitic origins to BAs (Figures 4B, 4C and 4F). Pseudotime ranking combined with k-means clustering (k = 5) of differentially expressed genes revealed that both lineages progress through an ordered five-stage developmental sequence toward BA fate (Figures 4G and S6A). GO analysis on stage-enriched genes identified several lineage-specific developmental pathways. For example, in endotome lineage at stages 2-5, pathways relating to VE and vSMC development were prominently enriched, including GO terms ‘smooth muscle cell differentiation’ and ‘regulation of hemopoiesis’ (Figure S6A). By contrast, the dermomyotome lineage at the corresponding stages showed preferential enrichment for SKM formation related GO terms (Figure S6A). Importantly, GO term ‘canonical Wnt signaling pathway’ was uniquely enriched in stage 3 of endotome lineage, corroborating the critical role of WNT activation between days 8-12 in promoting endotome-derived BA formation (Figures 3H-3K and S6A). Beyond these pathway-level distinctions, our extracted trajectory analysis resolved early BA precursors (stages 2-3, corresponding to endotome-BA lineage cultures at days 8-12) within endotome lineage that are characterized by VE and vSMC progenitor markers (e.g., *PENK*, *PLVAP* and *DLX1*),^43,72,73^ whereas dermomyotome-derived BA progressed through an *EN1+*/*GATA6*+ precursor stage (Figure 4H).^10,34,51^

To benchmark the two origin-specific BA populations and cross-validate their identities, we merged the BA clusters from the two lineages. Endotome-derived BAs exhibited elevated expression levels of WNT activators (*RSPO3*, *VWCE, NR4A1*) and effectors (*CTNNB1*, *TCF7L2*) compared with dermomyotome-derived BAs (Figure 4I), indicating sustained Wnt signaling activity specifically in the endotome-derived BA lineage. Furthermore, differentially expressed genes in endotome-BAs were enriched for cell proliferation and negative regulation of stress-induced kinase cascade (Figure S6B), hinting at a potential resistance to cellular stress and aging, consistent with prior observations in mice.^16^ To contextualize our cultured cells within established adipocyte landscape, we integrated them with scRNA-seq datasets from *in vitro*-derived beige adipocytes (this study) and *in vivo* adipose tissues (GSE164528, GSE185623 and GSE176171).^10,43,74^ The list of expressed genes in mouse datasets was converted to their human orthologs (Figure S6C). Hierarchical clustering revealed that endotome-derived BAs grouped closely with human adult and perinatal mouse peri-aortic BATs, whereas our dermomyotome-derived BAs clustered with mouse interscapular BAT (Figure 4J), recapitulating the expected in vivo relationships.^10,43,56^ Side-by-side comparison of BA differentiation markers further confirmed that both cultured BA populations reproduced the expression signatures of their respective *in vivo* tissue counterparts (Figure S6D). Interestingly, MSC-derived beige adipocytes displayed transcriptomic similarities with most included datasets, reflecting the ambiguous molecular distinction between beige and brown adipocytes.^5,59^ Collectively, these comparative analyses establish that peri-aortic and interscapular BAs follow parallel yet molecularly distinct developmental sequences, resulting in terminal BA populations that recapitulate the transcriptomic features of their respective anatomical depots (Figure 4K). Briefly, endotome-derived BA specification proceeds directly through a VE/vSMC-competent intermediate without overt SKM commitment, whereas dermomyotome-derived BA genesis requires a protracted SKM maturation phase followed by a GATA6+ precursor transition.

### BAs of endotome are thermogenic and metabolically active

Generally, BAs are characterized by their metabolic potential to generate heat in response to a β-adrenergic stimulus and downstream activation of the cAMP pathway, a physiological process that can be mimicked by treatment with forskolin (FSK) in cultures.^3,75^ To characterize the thermogenesis capacity of peri-aortic-like BAs of endotome, we measured their oxygen consumption rates (OCR) and extracellular acidification rates (ECAR) using a Seahorse metabolic flux analyzer. OCR and ECAR reflect oxidative phosphorylation and glycolytic flux, respectively. Using our previously established white adipocyte (WA) cultures as control,^3,56^ day-33 BA cultures exhibited marked increases of OCR and ECAR, which were further enhanced by a 24-hour FSK pre-treatment (Figure 5A-5E). WAs showed minimal responsiveness to FSK, as expected.

**Figure 5.**
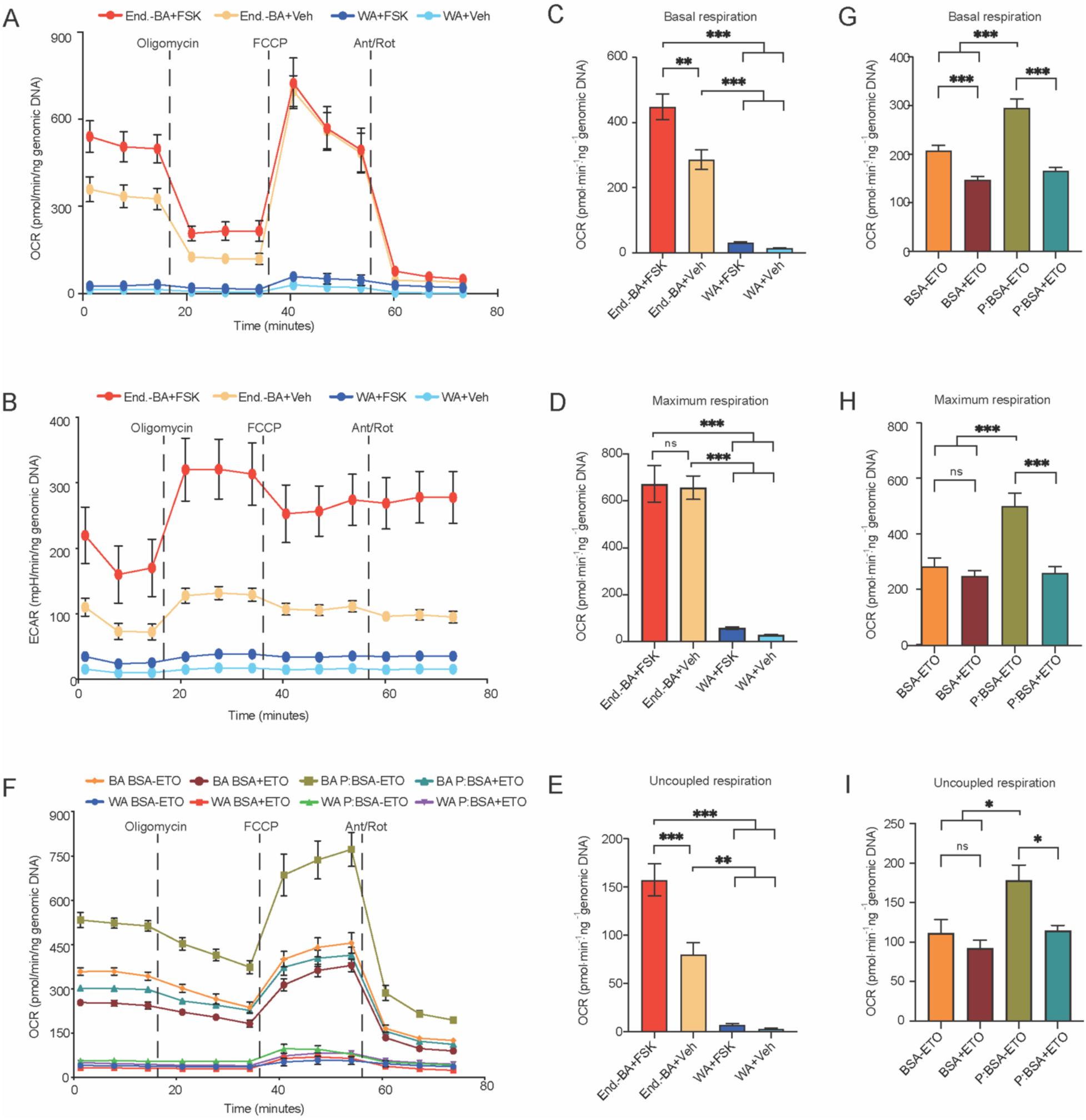
Functional analysis of endotome-derived Bas. (A) Oxygen consumption rates (OCRs) of endotome-derived BAs and MSC-derived white adipocytes (WAs) measured by Seahorse flux assay. Cells were treated with 10 µM Forskolin (FSK) or DMSO vehicle (Veh), 24 hours prior to the assay. 2 µM Oligomycin, 2 µM FCCP, and 5 µM antimycin A/rotenone (Ant/Rot) were injected at the indicated time points. n = 8. Raw readouts were normalized to genomic content per well. (B) Extracellular acidification rates (ECARs) of corresponding conditions in (A). (C-E) Quantification of basal (C), maximal (D) and uncoupled (E) respiration rates measured from the OCR assay in (A). (F) OCR measurements of BAs and WAs showing the capability of extracellular fatty acid oxidization using palmitate as a metabolic substrate. Cells were treated with 150 µM conjugated palmitate:BSA (P:BSA) or BSA alone, with or without 4 µM Etomoxir (ETO). n = 8 for WAs and n = 16 for BAs. (G-I) Quantification of basal (G), maximal (H) and uncoupled (I) respiration rates measured from the OCR assay in (F). Data were presented as mean ± SEM. ns, no statistical significance. *p < 0.05, **p < 0.01, ***p < 0.001

Another key feature of functional BAs is their ability to oxidize fatty acids for non-shivering thermogenesis.^3,76^ To assess this ability in endotome-derived BAs, we measured their OCR shifts in the presence or absence of palmitate (BSA alone as control) after a 24-hour starvation. As expected, palmitate treated endotome-derived BAs showed elevated basal and maximal respiration (Figures 5F-5H), whereas WAs responded poorly (Figure 5F). Etomoxir (ETO), an inhibitor of carnitine palmitoytransferase-1, reduced OCRs in these BAs (Figures 5G-5I). Consistent with a BA identity as previously described,^3^ endotome-derived BAs in this study underwent uncoupled respiration in response to FSK and utilized palmitate (Figures 5E and 5I) In summary, these experiments demonstrated that *in vitro* peri-aortic-like BAs of endotome origin exhibit elevated rates of uncoupled respiration that can be stimulated by pro-thermogenesis cues, as in *in vivo*.

### Endotome cells are multipotent that can give rise to vSMC and VE *in vitro*

BAT vasculature is critical for maintaining the high metabolic rates of BA and keeps BAs from ‘whitening’.^77–79^ After birth, aortic PVAT harbors various progenitors of vSMC and VE populations.^13,14,23,25,43^ Notably, peri-aortic BAs are reported to be more resistant to ‘whitening’ than interscapular BAs.^15,16^ Because endotome cells expressed vSMC and VE lineage tracing genes such as *PENK*, *TAGLN* and *PLVAP* (Figure 3D), we sought to assess whether endotome can directly give rise to vSMC and VE cell types, which would expand its utility for reconstituting the vasculature of peri-aortic BAT.

We first demonstrated the differentiation potential of day-8 endotome cells to become SMCs under treatment of 10 ng/ml HGF and 2 ng/ml IGF-1 (Figure 6A).^71,80^ Between days 11 and 13, marker genes for vSMC (*ACTA2*, *CNN1*) and contractile SMC (*MYOCD*, *MYH11*) significantly increased, at the expense of endotome identity (*EBF2*) (Figure 6B). Immunostaining of smooth muscle heavy chain 11 (MYH11), calponin 1 (CNN1) and α-smooth muscle actin (ACTA2) confirmed the contractile vSMC identity at day 13 (Figure 6C).^81^ To assert the direct contribution of endotome cells to SMCs, we performed a short-term lineage labeling experiment to trace EBF2-mScarlet-expressing cells to SMCs at days 9-11, using the EBF2-mScarlet H9-hESC line (Figure 6D).^18,82^ We took advantage of the perdurance of mScarlet to label cells that expressed EBF2 prior to the expression of calponin 1. At day 9, EBF2 and mScarlet proteins co-localized in the differentiating cells before the emergence of calponin 1. At day 11, endogenous EBF2 protein was undetectable, while the anti-RFP antibody still recognized the retained mScarlet protein, overlapping with the emergence of calponin 1, demonstrating the direct lineage contribution of EBF2-expressing endotome to a vSMC fate. Subsequent scRNA-seq analysis of day-5 (14,857 cells) and day-12 (9,421 cells) SMC cultures revealed two major populations of endotome-derived vascular SMCs, synthetic vascular SMC (syn-vSMC and contractile vascular SMC (con-vSMC) (Figure 6E). Angueira et al. identified several distinct SMC populations from aortic PVAT in postnatal mice, with a gradient of BA genesis competency.^43^ Interestingly, *MYH11*, *PRDM16*, *CD81*, *CD55*, *COX5A* and *COX7A2*, which were identified as SMC-related BA progenitor markers in the Angueria report, were also differentially expressed in our syn-and con-vSMC cultures at days 13-20 (Figure 6E). However, other important BA related markers such as *DPP4*, *PPARG* and *FABP4,* were absent from our SMC cultures (Figure 6E).^43,83^ Further studies are needed to determine if these SMC populations are brown adipogenic.

**Figure 6.**
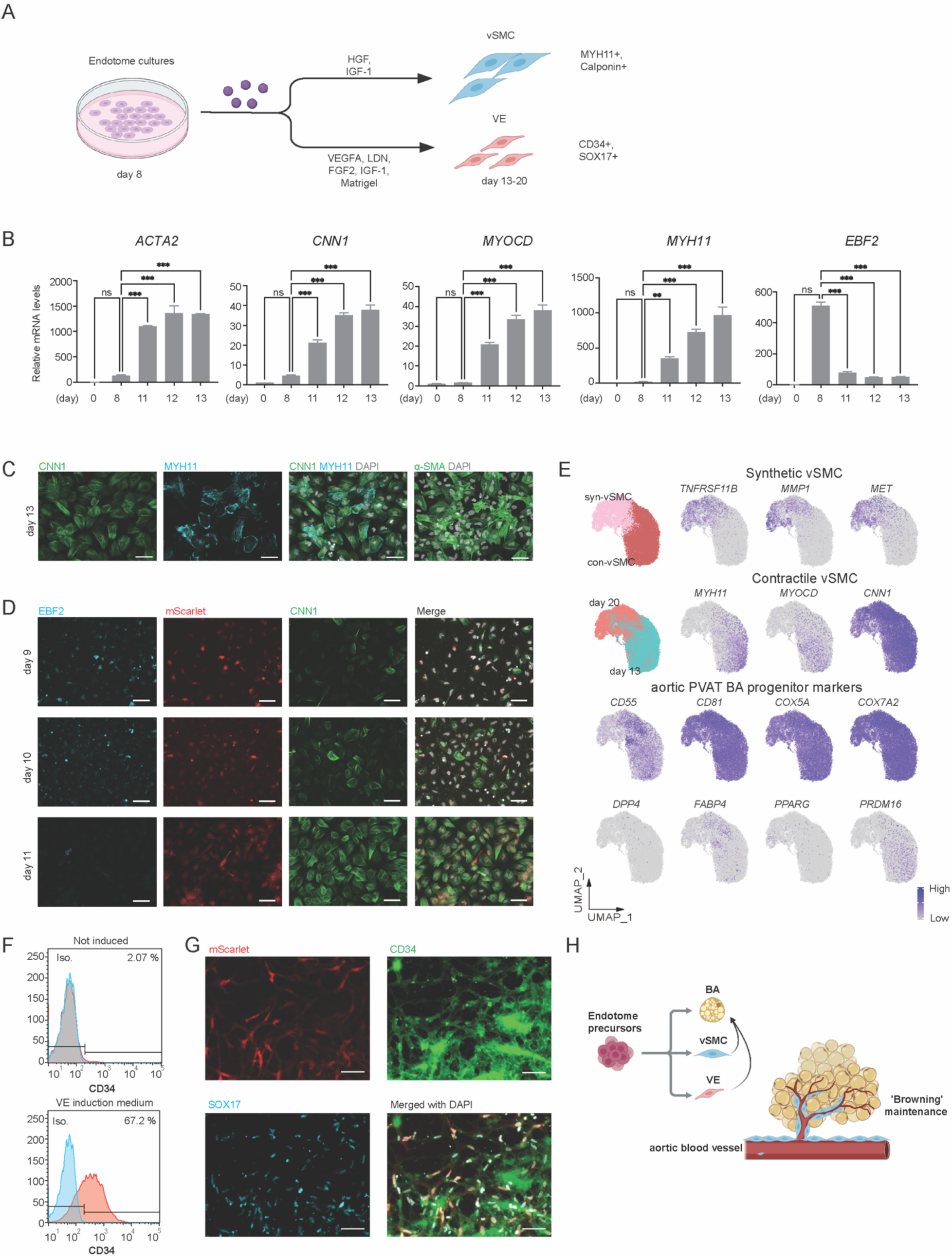
Endotome precursors are multi-potent and can give rise to vascular smooth muscle (vSMC) and vascular endothelial (VE) descendants. (A) Schematic illustrating the generation of vSMC and VE from endotome cells. (B) Real-time qPCR results showing the gradual increases of vSMC marker genes at indicated time points. n = 3. Data were presented as mean ± SEM. ***p < 0.001 (C) Immunostaining with antibodies against Calponin (CNN1), Myosin heavy chain 7 (MYH7) and alpha-smooth muscle actin (α-SMA, encoded by *ACTA2*) on day 13 in endotome-derived vSMC cultures. n > 3. Scale bars, 100 µm. (D) Immunostaining with antibodies against EBF2, mScarlet and Calponin (CNN1), showing the tracing of EBF2-mScarlet-expressing cells toward CNN1-expressing cells during days 9-11. (E) UMAP projection of integrated endotome-derived vSMC cultures (days 13 and 20), with featureplots showing marker genes for vSMC subtypes and BA progenitors. (F) Flow cytometry analysis of fraction of CD34 positive VE cells derived from endotome cultures. (G) Immunostaining with antibodies against mScarlet, CD34 and SOX17 in VE cultures, showing the tracing of EBF2-mScarlet-expressing endotome cells toward CD34/SOX17-expressing VE fate on day 15. n > 3. Scale bars, 100 µm. (H) Schematic summary of the multi-lineage potential of endotome cells in reconstituting the vasculature in peri-aortic BA tissue.

We next tested if VE cells can also be induced from endotome. Given that endotome resides within the lateral somite signaling environment, where VEGF signaling activation and BMP inhibition promote VE differentiation,^20,21,47,84,85^ we developed a protocol using 10 ng/ml VEGFA, 500 nM LDN, 20 ng/ml FGF2, 2 ng/ml IGF-1 and 1% Matrigel (Figure 6A). Culturing day-8 endotome cells under this condition for 7 days induced 67% CD34+ VE cells (Figure 6G), in contrast to the blank control treated with FGF2 and IGF-1 alone. Similarly, indirect lineage tracking using the EBF2-mScarlet line further confirmed the differentiation of EBF2-expressing endotome cells to a CD34+/SOX17+ VE identity on day 15 (Figure 6H). In summary, these data established that *in vitro*-derived endotome cells are vasculogenic that may be functionally analogous to the progenitors of vSMC and VE, which constituting the supporting tissues in aortic PVAT (Figure 6I).^59,86^

## Discussion

In this study, we establish the EBF2+ endotome as an early-specified, MYF5-independent somite compartment that gives rise to peri-aortic BAs. Peri-aortic BAT represents a distinct thermogenic depot, conserved in perinatal mice and adult humans, that shares core functional attributes with interscapular BAT, including dependency on the BA transcription factor EBF2, high levels of UCP1 expression and uncoupled respiration (Figures S2E, 2H-2J and 5). To characterize the molecular identity and ontogeny of this depot, we interrogated single cell profiling of mouse organogenesis alongside human trunk embryoid models. This approach reveals that endotome represents the *PAX3*+/*MYF5*- somitic origin of BAs and are molecularly primed by a TGF-β-induced EMT program. Recapitulating the sequential signaling cues of TGF-β activation, BMP inhibition and subsequent Wnt activation enables efficient BA commitment, yielding cells that transcriptionally and functionally resemble *in vivo* peri-aortic BAT.

Our data position the endotome as the earliest derivative of the *PAX3*+ somite, with *EBF2* detection as early as E7.75 in mice and day 6 in hTEM.v3, preceding the onset of *MYF5*⁺ dermomyotome specification (E8.5 and day 7, respectively) by approximately 12-24 hours. We posited that this spontaneous endotome differentiation coincides with the emergence of somite-originated VE in various mouse and human trunk organoid models,^18,31,87^ where *PAX3*+ pan-somite cells dominate without detectable dermomyotome or sclerotome derivatives. This suggests that endotome is an autonomously specified vascular-associated compartment. Independent lineage tracing using *Tagln*-Cre further supports this, showing that early *Tagln*+ cells (E8.5 onward) contribute to perivascular structures, including aortic SMCs and adipogenic fibroblasts, and ultimately to thermogenic adipocytes.^28^ Notably, our endotome cultures exhibit elevated *TAGLN* expression at days 8-12, corresponding to BA fate commitment stage (Figures S5B and S5E). Therefore, direct lineage-tracing and functional perturbation studies will be needed to establish whether the reported early *Tagln*+ BA precursros represent the endotome proper *in vivo*.

Despite this early molecular specification, peri-aortic BAT depots only become structurally organized during the perinatal period (E18-P3) in mice, coincident with a burst of vasculogenesis.^43^ This temporal gap (roughly 10 embryonic days between lineage commitment and tissue organization) suggests that endotome precursors remain poised until the perinatal niche trigger their terminal differentiation. This unique developmental dynamic markedly contrasts with the formation of interscapular BAT, where *Pax3+*/*Myf5+*/*Pax7*+ dermomyotome precursors continuously generate BA progenitors and BAs throughout mid-gestation (E13.5-E15.5).^8–10^ Notably, this developmental dichotomy is reflected by our concurrent human BA differentiation cultures from the two somitic lineages *in vitro*. Endotome-derived BA formation requires only 12 days post-induction, whereas dermomyotome-derived BA formation requires 24 days (Figure 4A and S5A-S5D). We attribute this accelerated BA formation to the intrinsically proliferative state of endotome-derived BAs, a distinct feature supported by GO enrichment analysis (Figure S6B). Given that our endotome-derived cultures require ∼45% less differentiation time and contain markedly fewer (∼50% less) non-target cell populations than their dermomyotome-derived counterparts (Figures 4A-4C),^10^ this protocol offers a more practical and scalable approach for *in vitro* BA generation.

Mechanistically, our study uncovers a signaling logic for the endotome that is fundamentally different from that in the dermomyotome lineage. In the report by Rao et al., 2023^10^, BMP7 and Wnt antagonist (C59) were used in combination with an adipogenic cocktail on day-20 EBF2+ precursors to induce GATA6+ BA precursors and subsequent interscapular BAs. To side-by-side compare endotome- and dermomyotome-derived BAs (Figures 4A and 4J), we employed the same BA induction medium containing TGF-β signaling inhibitor (SB-431542),^3,50^ but omitted exogenous BMP7 and C59, in which GATA6+ precursors still emerged in our adopted dermomyotome-BA cultures as expected. Despite these technical differences, the conceptual divergence is obvious. Whereas Rao et al., 2023 study concluded that interscapular BA development requires BMP activation and Wnt inhibition, we demonstrated that the EBF2+ endotome cells require precisely the opposite signaling activities for BA fate specification. BMP4 suppresses EBF2 expression levels (Figures S2E and S2F) and BMPs (BMP4/5/7) in general abolish BA genesis potency of endotome cells (Figures 2H-2K). The requirement for exogenous Wnt activation could be rationalized by our CUT&Tag data. Although EBF2 binds to putative enhancers at genes mediating TGF-β signaling and EMT processes (Figures 3F and S3E), it does not occupy core Wnt pathway genes, such as *CTNNB1*, *TCF7*, *TCF7L2* and *NR4A1* (data not shown), thus necessitating a pulse of Wnt activation for full BA commitment (Figures 3H-3K). This Wnt dependency agrees with the recent finding by Loureiro et al., 2023 that a transient perturbation of Wnt signaling is sufficient to cause long-term changes in adipogenic program within the mesenchymal stromal niche.^60^ Our mechanistic framework therefore will help understand stage- and depot-specific BA development, particularly within peri-aortic BAT and PVAT depots. Notably, while Wnt signaling has been linked to interscapular BA ‘whitening’ and type 2 diabetes,^70,88–90^ sustained Wnt signaling activity has been observed in our endotome-derived BAs (Figure 4I). This leads to a pivotal open question: whether unique feature of peri-aortic BAs can provide a sustainable and ‘whitening’-resistant source of BAs for cell-based regenerative medicine.^16^

Additionally, we observed increased TGF-β priming upregulated both transcript and protein levels of SOX9 (Figures 2D, 2E and 3D). Similar to our findings, Sox9 was recently identified as a combinatorial marker alongside Gata6 and Ebf2 for non-interscapular BA origins in mice.^42^ In that study, the authors demonstrated that Sox9 transiently labeled BA precursors within cervical BAT (E9.5-E15.5) and scapular BAT (E10.5-E15.5) depots. Given that endotome emerged at E7.75-E8.5 (Figures 1A-1C) and that peri-aortic BAT forms at E18-P3,^19,43^ we proposed that a combinatorial lineage-tracing strategy incorporating Ebf2+/Sox9+/Gata6- markers could be effectively employed to label endotome-derived peri-aortic population in mice. Uncovering the developmental logic and regulatory networks governing this previously overlooked endotome-derived BAT depot bears profound implications for both disease pathophysiology and regenerative medicine. Given BAT’s important role in energy expenditure and cardiometabolic health, elucidating the distinct molecular identity and commitment cues of this aortic-adjacent lineage may unveil novel diagnostic biomarkers and pharmacological targets for obesity, insulin resistance, and associated cardiovascular complications.

For proof-of-concept that endotome constitutes a self-sufficient, all-in-one source for reconstituting vasculature of BAT, we demonstrated that day-8 endotome cells directly give rise to vSMC and VE populations (Figure 6), primarily in response to HGF and VEGFA, respectively. This differentiation capacity is functionally consequential, given the growing evidence implicating the aortic PVAT niche as a key modulator of BA expansion, homeostasis, metabolism and ‘browning’ maintainence.^43,77,78,91–93^ Clinically, thoracic PVAT composed of brown adipose tissue in the healthy state and undergo a brown-to-white transition, which was recognized as a driving factor in cardiovascular disease and obesity. Accordingly, the ability to derive both vSMC and VE lineages from the same endotomal origin as peri-aortic BA precursors offers a unique experimental toolkit for the *in vitro* assembly of ontogeny-matched, functional peri-aortic-like BAT vasculature (Figure 6H). Moreover, the restricted temporal window and specific anatomical niche of these precursors/progenitors present a versatile strategy for regenerative interventions. Harnessing or selectively reactivating this endogenous precursor/progenitor pool could shed light on BA organoid-based therapies and tissue-engineering approaches aimed at restoring functional BA vasculature in metabolic disorders or age-related thermogenic decline.^5,86^ Collectively, our findings could therefore help dissect the vascular-adipose interplay for therapy strategies involving the transplantation of human BAs generated *in vitro* for treatment of obesity and related metabolic disorders.

### Limitations of the study

First, the transition from endotome-derived vSMC to mature adipocytes remains incompletely defined. Angueira et al. 2021 demonstrated that adipogenic SMCs (*Myh11+, Pdgfra*-*, Pparg+)* contribute to aortic PVAT formation.^43^ However, whether our *MYH11*+/*ACTA2*+/*PPARG*- vSMCs can differentiate into *HIC1*+/*PDGFRA*+ adipogenic fibroblasts or *PPARG*+ BA progenitors remains unresolved. We did not perform direct BA induction from endotome-derived vSMCs, and aortic PVAT contains distinct fibroblast populations with adipogenic potential separate from the vSMC lineage.^43^ Definitive lineage-tracing and clonal analyses are required to determine if the *MYF11*+/*ACTA2*+ cells of *in vitro* endotome give rise to BAs via a transitional stage or represent a non-adipogenic parallel population, which is critical for evaluating their utility as a renewable source of BA precursors.

Second, the contribution of endotome to *in vivo* peri-aortic BAT was primarily inferred through transcriptional similarities and lineage-tracing implications in literatures rather than direct spatiotemporal fate mapping in humans, which is inherently unattainable. Although our previous hTEM platform mirrors early mouse developmental sequences (E7.5-E8.5),^18^ future humanized xenotransplantation or organoid co-culture models will be necessary to definitively confirm the functional equivalence of endotome-derived BAs to their *in vivo* counterparts at postnatal stages. Addressing these limitations will be crucial for translating this platform toward therapeutic applications.

## Supporting information

Key Resources Table

## Resource availability

The raw and processed sequencing data are available in the GEO repository GSE336917 with a reviewer’s token sfijckqejbivnah. Source code for analyzing the sequencing data is available at: https://github.com/yhao10569/YH_brown_adipocytes_2026

## Acknowledgement

We thank the Core Laboratories of School of Biomedical Sciences at the Chinese University of Hong Kong for the service of single cell sorting, histology sections and imaging platform. This study was supported by grants to S.D. from the Research Grants Council of Hong Kong (General Research Fund) and the Hong Kong Jockey Club Charities Trust. S.D. is a Global Stem Scholar and Director of the JC STEM Lab of Stem Cells and Regenerative Medicine.

## Author contributions

Conceptualization, T.-M.W. and S.D.; methodology, T.-M.W. and H.Y.; data collection, H.Y., T.-M.W., W.-M.X., K.-X.T., E.S.-K.N., A.Y.-F.K.; software, H.Y., T.-M.W., W.-M.X., K.-X.T., S. P.; writing - draft, T.-M.W.; writing - editing & revision, T.-M.W., H.Y., S.D.; supervision, T.-M.W. and S.D.; funding acquisition, S.D.

---

**Figure S1.**
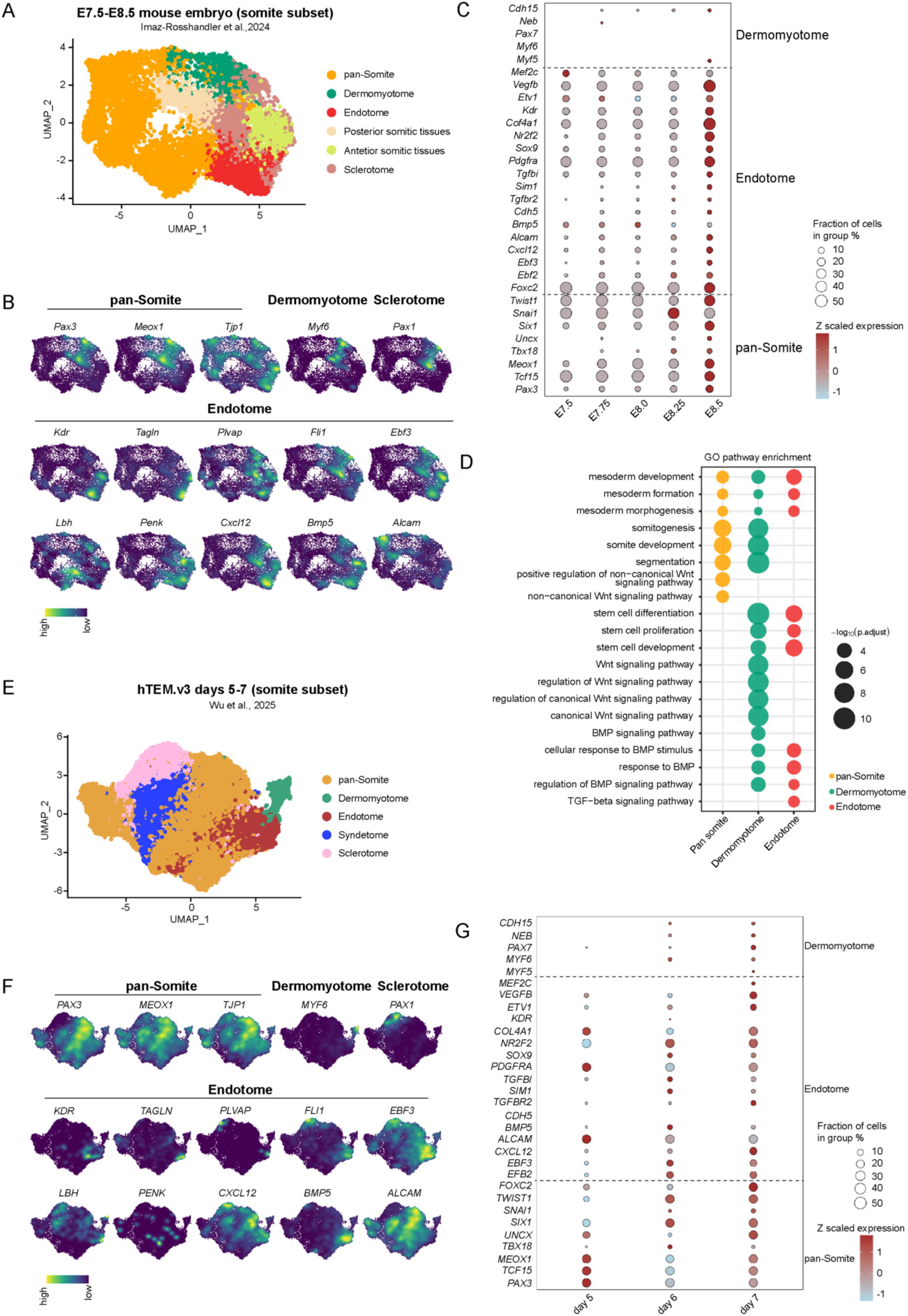
Embryonic development of endotome and dermomyotome in somites. Related to Figure 1 (A) UMAP showing cells represneting somite development from E7.5-E8.5 mouse embryos in Imaz-Rosshandler et al., 2024 study. (B) UMAP showing expression levels of a curated list of cluster-specific genes in mouse somite compartments. UMAP embeddings are corresponding to (A). (C) Dotplot showing scaled expression of extended marker genes over time. Transcripts in cells were extracted from (A). Fraction of detected cells below 1% were omitted. (D) Gene ontology analysis of top 100 differentially expressed genes from indicated cell types. (E) UMAP showing cells representing somite development from human trunk embryoid model version 3 (hTEM.v3; days 5-7) in Wu et al., 2025 study. (F) UMAP showing expression levels of a curated list of cluster-specific genes in human somite compartments. UMAP embeddings are corresponding to (E). (G) Dotplot showing scaled expression of extended marker genes over time. Transcripts in cells were extracted from (E). Fraction of detected cells below 1% were omitted.

**Figure S2.**
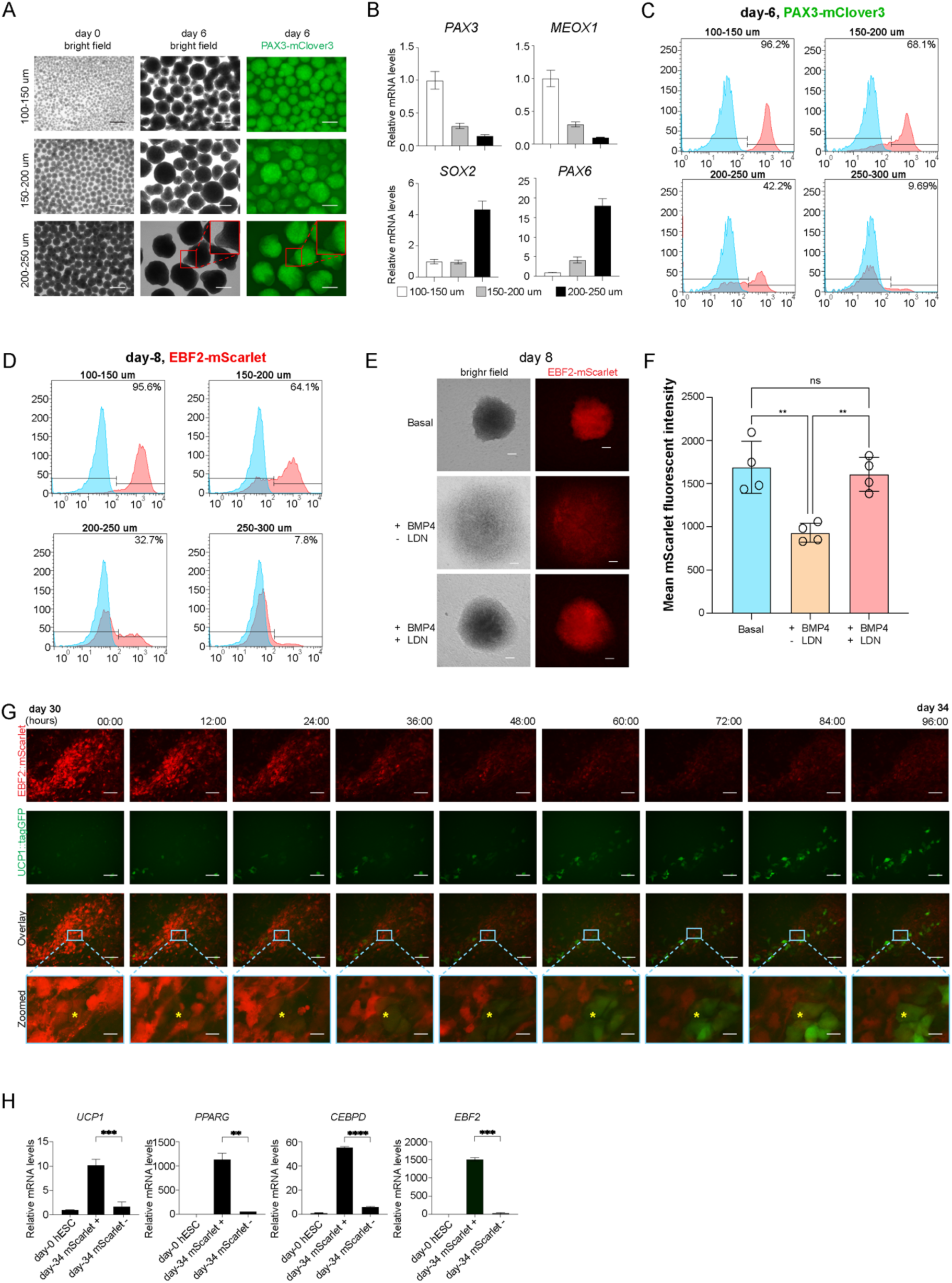
De novo differentiation of PAX3+ somite, EBF2+ endotome cells, and subsequent UCP1+ Bas. Related to Figure 2 (A) Representative images showing PAX3-mClover3 expression in size-controlled hESC spheroids. Red boxes highlighted the heterogenous PAX3-mClover3 expression observed on day 6 in larger (200-250 µm) hESC spheroids. Scale bars, 100 µm. (B) Real-time qPCR analysis of *PPARG*, *ADIPOQ* and *UCP1* on day-6 somite cultures in (A). Data were presented as mean ± SEM. n = 3. (C) Flow cytometry analysis showing fraction of PAX3-mClover3 positive cells in day-6 somite cultures derived from (A). Reproduced from over three biological batches. (D) Flow cytometry analysis showing fraction of EBF2-mScarlet positive cells in day-8 endotome cultures (without TGF-β1) derived from (A). Reproduced from over three biological batches. (E) Fluorescence images of EBF2-mScarlet signals in day-8 endotome cultures under three treatment conditions: basal medium containing 0.5 µM LDN without TGF-β1, basal medium supplemented with 10 ng/ml BMP4 alone, and basal medium supplemented with 10 ng/ml BMP4 and 0.5 µM LDN. Scale bars, 100 µm. (F) Flow cytometry quantification of EBF2-mScarlet intensities from (E). n = 4. Data were presented as mean ± SEM. ns, no statistical significance. **p < 0.01. (G) Time-lapsed imaging of days 30-34 BA cultures derived from day-8 endotome cells (without TGF-β1). Yellow stars tracked the continuous transition from EBF2-expressing BA progenitors to UCP1-expressing BAs. Scale bars, 100 µm. (H) Real-time qPCR analysis of BA progenitor markers in day-34 BA cultures from (G). To exclude lipid-rich BAs, the cultures were filtered through a 35 µm strainer prior to RNA extraction. n = 3. Data were presented as mean ± SEM. **p < 0.01, ***p < 0.001.

**Figure S3.**
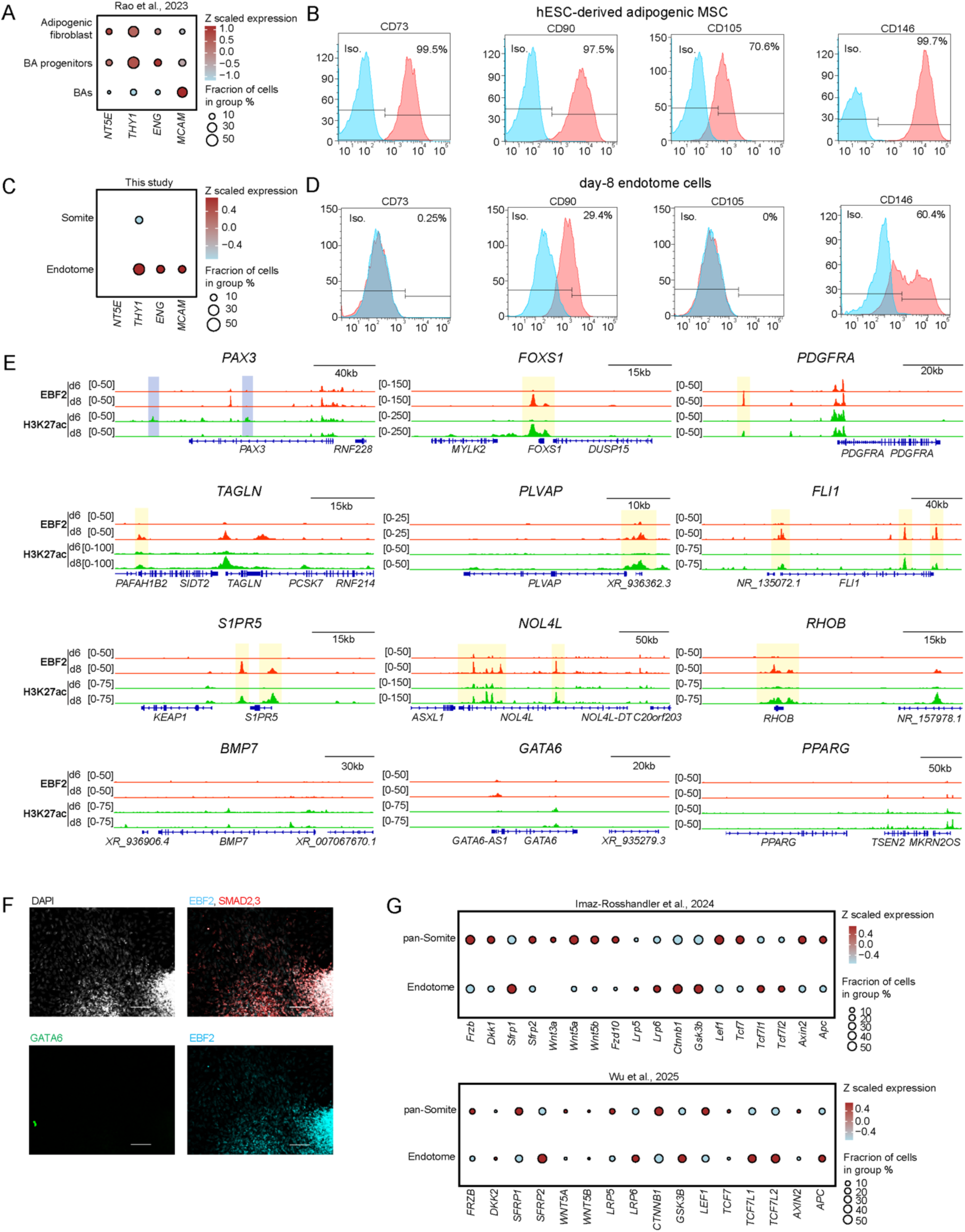
Molecular profiling of TGF-β1-stimulated endotome cells. Related to Figure 3 (A) Dotplot showing scaled expression of adipogenic surface marker genes including *NT5E* (CD73), *THY1* (CD90), *ENG* (CD105) and *MCAM* (CD146) in indicated hiPSC-derived BA populations in Rao et al., 2023 study. (B) Flow cytometry analysis of adipogenic surface markers of CD73, CD90, CD105 and CD146 for hESC-derived MSCs in this study. (C) Dotplot showing scaled expression of adipogenic surface marker genes including *NT5E* (CD73), *THY1* (CD90), *ENG* (CD105) and *MCAM* (CD146) in day-6 somite and day-8 endotome cells. Cells analyzed were from Figure 3A. Fraction of detected cells below 1% were omitted. (D) Flow cytometry analysis of adipogenic surface markers of CD73, CD90, CD105 and CD146 in day-8 TGF-β1-stimulated endotome cells. (E) Genome browser tracks showing CUT&Tag signals for EBF2 and H3K27ac (putative enhancer marker) in day-6 somite and day-8 endotome cells. Shaded areas in blue highlighted the putative enhancers in somite cells. Yellow shades highlighted EBF2-bound putative enhancers in endotome cells. (F) Immunostaining of day-8 TGF-β1-stimulated endotome cells using antibodies against EBF2, SMAD2,3 and GATA6. Scale bars, 100 µm. (G) Dotplot showing scaled expression of core WNT pathway genes in pan-somite and endotome cells from (top) mouse E7.5-E8.5 in Imaz-Rosshandler et al., 2024 and (bottom) human trunk embryoid model in Wu et al., 2025. Fraction of detected cells below 1% were omitted.

**Figure S4.**
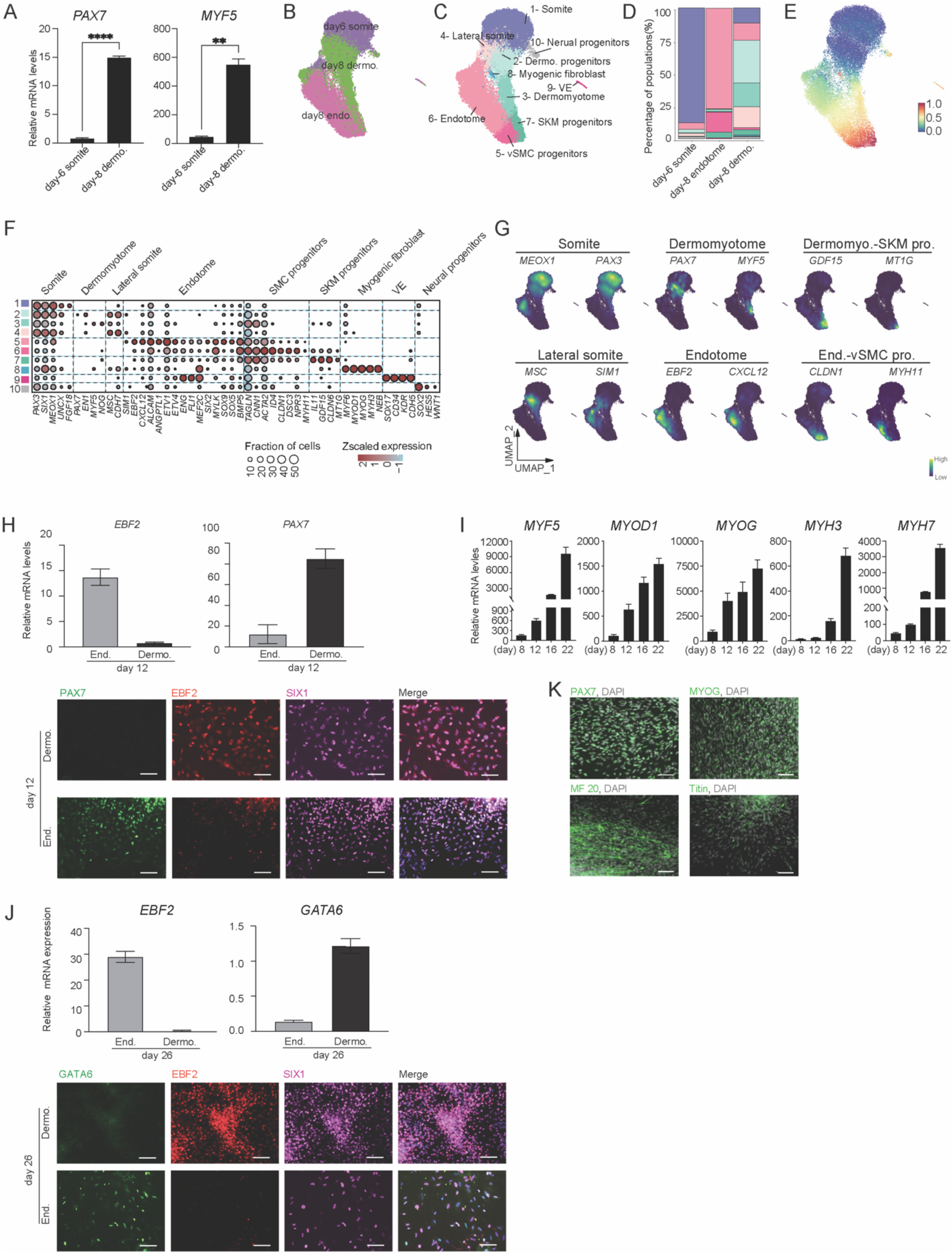
Endotome precursors undergo a MYF5-/PAX7-/GATA6- developmental path toward BA fate. Related to Figure 4 (A) Real-time qPCR results showing the expression of dermomyotome markers in day-8 dermomyotome cultures. n = 3. Data were presented as mean ± SEM. **p < 0.01, ****p < 0.0001. (B) UMAP showing the integration of day-6 somite, day-8 endotome and day-8 dermomyotome cells. (C) UMAP embedding displaying clusters of identified cell types in (B). (D) Stack plot showing fractions of identified cell types in (C). (E) UMAP embedding of pseudotime scores, inferring the developmental progression of cells shown in (D). (F) Dotplot showing scaled expression of an extended list of marker genes in identified cell types in (D). Fraction of detected cells below 1% were omitted. (G) Featureplots showing representative lineage-specific marker genes on UMAP trajectory shown in (D). (H) qPCR (top) and immunostaining (bottom) results showing that PAX7 and EBF2 were differentially expressed in day-12 cultures of endotome and dermomyotome cells. SIX1, somitic lineage marker. Scale bars, 100 µm. (I) qPCR measurements of dermomyotome-derived skeletal muscle marker genes at day 8, 12, 16 and 22. (J) qPCR (top) and immunostaining (bottom) results showing that GATA6 and EBF2 were differentially expressed in day-26 cultures of endotome and dermomyotome cells. SIX1, somitic lineage marker. Scale bars, 100 µm. (K) Immunostaining with antibodies against PAX7, Myogenin (MYOG), Scaromeric myosin heavy chain (MF 20) and Titin on day 26 in dermomyotome lineage cultures. Scale bars, 100 µm.

**Figure S5.**
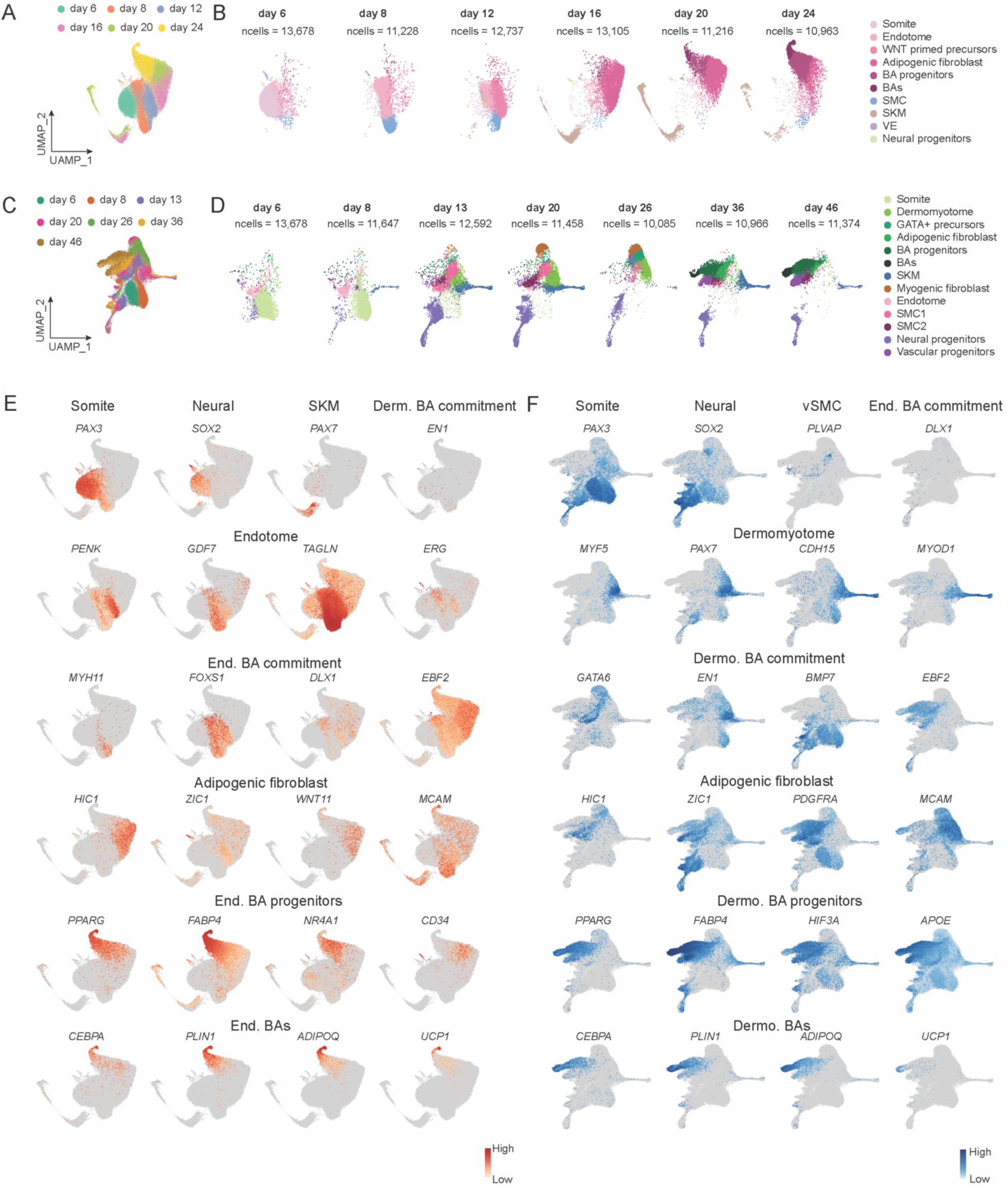
Endotome and dermomyotome exhibit lineage-specific molecular markers during BA fate specification. Related to Figure 4 (A) UMAP showing the integrated cells of *in vitro* endotome-derived BA cultures at days 6, 8, 12, 16, 20 and 24. (B) UMAP embedding showing the identified cell types on the indicated day, with colors and annotations as in Figure 4B. (C) UMAP showing the integrated cells of *in vitro* dermomyotome-derived BA cultures at days 6, 8, 13, 16, 20,26, 36 and 46. (D) UMAP embedding showing the identified cell types on the indicated day, with colors and annotations as in Figure 4C. (E and F) Featureplots showing a curated list of lineage-specific and BA-staged marker genes, shown across in time-integrated datasets derived from endotome (E) and dermomyotome (F).

**Figure S6.**
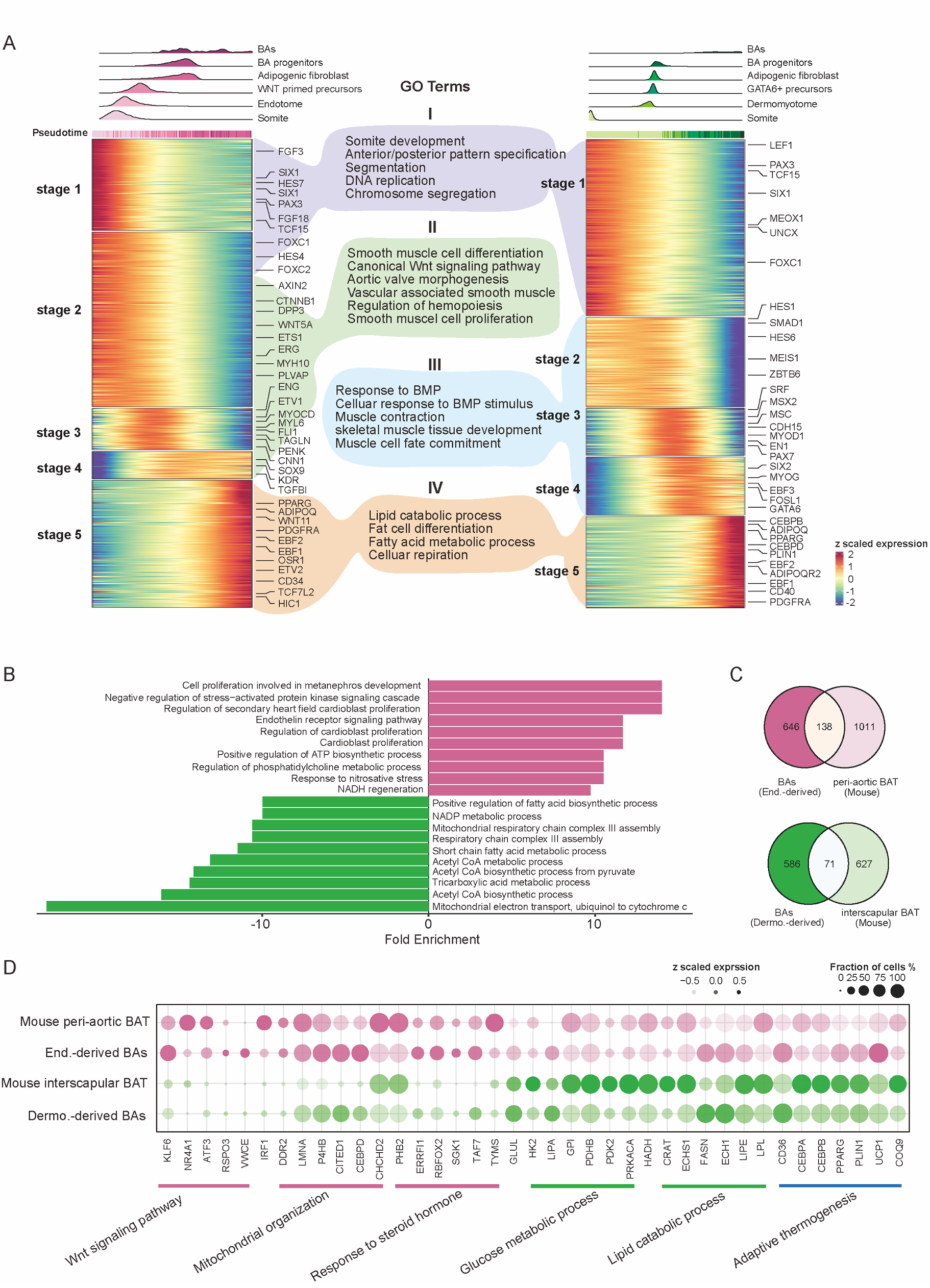
Endotome and dermomyotome undergo distinct biological processes toward BA fate. Related to Figure 4 (A) Top, ridge plot showing pseudotime-ranked cell populations depicting endotome-and dermomyotome-derived BA progressions, with colors and cell identities as in Figure 4F. Bottom, heatmap showing k means clustering (k = 5) of differentially expressed genes in corresponding pseudotime-ranked cells. Representative genes and Gene ontology (GO) terms were listed next to the clustering-defined developmental stages. GO analysis was performed using differentially expressed genes as input at each stage from respective lineage. (B) Barplot showing the top10 differentially enriched GO terms in endotome- and dermomyotome-derived BAs. (C) Venn diagram showing the conserved transcripts between our *in vitro*-derived human BAs and *in vivo* mouse BATs. Mouse peri-aortic BAT (GSE164528) and mouse interscapular BAT (GSE185623) datasets are publicly available. Expressed genes in mouse datasets were converted to their human orthologs. (D) Dotplot showing scaled expression of selected genes conserved between human and mouse BAs as in (C). Fraction of detected cells below 1% were omitted.

## Methods

### Cell lines and cultures

The following human ESC lines were used: H9-hESC (WiCell, WAe009-A) and derived lines of PAX3-mClover3/EBF2-mScarlet reporter,^18^ EBF2-mScarlet/UCP1-tagGFP reporter, TGFBR2-LOF (depletion of exon 5 and resultant coding frame shift). All cell lines were routinely tested by PCR, to ensure that they were not contaminated with mycoplasma.

The pluripotency of these hESC lines were maintained using in-house HAIF medium as described previously.^18,94^ Briefly, 5 × 10^4^/cm^2^ cells were seeded onto TC surface Easydish (Thermo, 150462) coated with 1:200 diluted Geltrex (Thermo, A1413302). When cells reach 80-90 % confluency, they were dissociated using Accutase (Thermo, 00-4555-56) and passaged using fresh HAIF medium containing 10 μM ROCKi for the first 24 hours. Medium was changed every 24 hours at 37 °C in 5% CO2.

The complete HAIF was reconstituted by supplementing the basic defined medium (DM) with 10 ng/mL human Heregulin β1 (Qkine, QK045), 10 ng/mL human Activin A (Qkine, QK001), 8 ng/mL human FGF2 G3 form (Qkine, QK052) and 200 ng/mL human IGF-1 LR3 form (Qkine, QK041). The basic DM was prepared by combing the following components, DMEM/F-12 (no glutamine) (Thermo, 21331020 or Servicebio, G4514), supplemented with 0.5% Probumin (Sigma, 821001), 1x Antibiotic-Antimycotic (Thermo, 15240062), 1x MEM NEAA (Thermo, 11140050), 1x Trace Elements A/B/C (Corning), 64 μg/mL Ascorbic acid magnesium (TCI, A2521), 10 μg/mL Transferrin (Athens Research and Technology), and 1 x GlutaMax (Thermo, 35050061), and then sterile-filtered using a vacuum filter unit (Thermo, 569-0020).

### Brown adipocyte differentiation from endotome and dermomyotome cells

BAs from the two lineages were both derived from PAX3+ somite cells using a modified protocol from Rao et al., 2023 and Wu et al., 2025,^10,18^ with somite specification efficiency being highest from smaller (100-150 μm in diameter) hESC spheroids (Figures S2A-S2C). To generate these spheroids, hESCs at 75-80% confluency were dissociated into single cells using Accutase. At day -2, 10^6^ cells were resuspended in 5 ml HAIF with 10 μM ROCKi into one well of an Ultra Low-attachment 6-well plate (Corning, 3471) and placed on an orbital shaker (Thermo, 88881104) at 130 rpm within a 37 °C, 5% CO_2_ incubator. After 24 hours, the medium was gently exchanged with fresh HAIF without ROCKi. At day 0, hESC spheroids with desired sizes were collected for somite specification. For larger spheroids, seeding densities (1-3 x 10^6^ cells per well) and shaking speed (110-130 rpm) were titrated. Larger cell number with slower agitation yielded in increased spheroid diameters, as shown in Figure S2A.

#### PAX3+ somtie induction

To induce PAX3+ somite formation, hESC spheroids were subjected to a two-step somitic mesoderm induction, transitioning from presomitic to somitic identity. At days 0-2, hESC spheroids were cultured in presomitic mesoderm induction medium, which was then replaced with somite induction medium for days 2-6. The plate was maintained on the orbital shaker at 130 rpm throughout somite formation period. The **presomitic mesoderm induction medium** consisted of DMEM/F12 (without glutamine) supplemented with 1x Insulin-Transferrin-Selenium (ITS; Thermo, 41400045), 3 μM CHIR-99021 (CHIR; MCE, HY-10182), 0.5 μM LDN-193189 (LDN; Tocris, 6053), 1x NEAA, 1x GlutaMAX and 1x Antibiotic-Antimycotic. The **somite induction medium** comprised of DMEM/F12 (no glutamine), 1x ITS, 1.5 μM CHIR, 0.5 μM LDN, 5% fetal bovine serum (FBS; Thermo, 30044333), 1x NEAA, 1x GlutaMAX and 1x Antibiotic-Antimycotic. 10 μM ROCKi was added for the first 24 hours, followed with daily medium changes.

#### BA differentiation of endotome lineage

To differentiate PAX3+ somites to endotome fate, day-6 PAX3+ somites were sedimented by gravity in a 50 ml conical tube, washed with endotome induction medium, and replated onto geltrex-coated TC surface multi-well plates (Thermo) with gentle agitation to ensure uniform distribution and prevent clumping. The **endotome induction medium** was made of DMEM/F12 (no glutamine), 15% KSR, 10 ng/ml human TGF-β1 (R&D, 7754-BH-005), 0.5 μM LDN, 2 ng/ml human IGF-1, 20 ng/ml FGF2, 2 ng/ml IGF-1, 1x NEAA, 1x GlutaMAX and 1x 1x Antibiotic-Antimycotic. 10 μM ROCKi was added for the first 24 hours, followed by daily medium changes to day 8.

At day 8, endotome cultures were dissociated using Accutase and passaged at density of 40,000 cells/cm^2^ on Geltrex-coated TC surface plates in WNT priming medium, followed with daily medium changes to day 12. The **WNT priming medium** comprised of DM supplemented with 3 μM CHIR, 20 ng/ml FGF2 and 1x ITS.

At day 12 onward, cells were cultured with BA induction medium, with medium change every other day, until the appearance of obvious lipid droplets (usually occurred after day 20). The **BA induction medium** consisted of DM supplemented with 2 μM rosiglitazone (Tocris, 5325), 1 μM dexamethasone (Tocris, 1126), 1 nM 3,3’,5-Triiodothyronine sodium salt (LT3; Aladdin L129594), 500 μM IBMX (Aladdin, I106812), 10 μM SB-431542 (Selleck, S1067), 200 ng/ml IGF-1 and 8 ng/ml FGF2.

#### BA differentiation of dermomyotome lineage

BA differentiation from dermomyotome was modified from Rao et al., 2023. The same replating procedure used for endotome induction was applied to somite spheroids at days 6-8, except that the medium was replaced with **dermomyotome induction medium**, which consisted of DMEM/F12 (no glutamine), 15% KSR, 3 μM CHIR, 10 ng/ml human HGF (PeproTech, 100-39), 2 ng/ml IGF-1, 1x NEAA, 1x GlutaMAX and 1x Antibiotic-Antimycotic.

At days 8-26, CHIR was omitted from the medium to allow the emergence of PAX7+ skeletal muscle progenitors and GATA6+ BA precursors (Figures S4H-S4K), as described in Rao et al., 2023. Medium was changed every other day. At day 26 onward, these dermomyotome-derived cells were cultured in BA induction medium, with medium changed every other day, until the appearance of obvious lipid droplets (usually occurred after day 36).

### Differentiation of vascular smooth muscle and endothelial cells from endotome

To direct lineage-specific differentiation, day-8 endotome cells were dissociated with Accutase, replated onto on Geltrex-coated TC surface, and cultured in either vascular smooth muscle (vSMC) induction medium or vascular endothelial (VE) induction medium. The **vSMC induction medium** consisted of DMEM/F12 (no glutamine), 15% KSR, 10 ng/ml human HGF (PeproTech, 100-39), 2 ng/ml IGF-1, 1x NEAA, 1x GlutaMAX and 1x Antibiotic-Antimycotic. The **VE induction medium** comprised DMEM/F12 (no glutamine), 10 ng/ml human VEGFA (Thermo, 100-20), 0.5 μM LDN, 2 ng/ml IGF-1 and 1% Matrigel (Mogengel, 827775). For each fate, differentiation markers were assessed at days 12-20.

### Differentiation of human beige and white adipocytes from hESCs

Beige and white adipocytes (WAs) were differentiated from hESC-derived adipogenic mesenchymal stem cells (MSCs) using a modified protocol from Singh et al., 2020.^56^

#### Generation of adipogenic MSCs

hESCs at 70% confluency (day 0) were cultured with STEMdiff-ACF mesenchymal induction medium (StemCell Technologies, 05240) for three days, with daily medium changes. From day 4 onward, medium was changed to MesenCult-ACF Plus medium (StemCell Technologies, 05445), also replenished daily, to promote MSC differentiation. Upon reaching 80% confluent, cells were passaged using ACF enzymatic disscoation solution (StemCell Technologies, 05426). By day 35, the successful generation of adipogenic MSCs was confirmed by flow cytometry analysis of surface markers including CD73, CD90, CD105 and CD146 (Figure S3B). The resulting adipogenic MSCs maintained stable expression of these surface markers for at least 10 passages, as verified by flow cytometry.

#### Beige adipocyte differentiation

For beige adipocyte differentiation, hESC-derived adipogenic MSCs were cultured in the B8 medium for 21 days, as described in Singh et al., 2020. The **B8 medium** consisted of DM supplemented with 200 ng/mL IGF-1, 8 ng/mL FGF2, 10 μM Y-27632, 2 μM rosiglitazone, 1 μM dexamethasone, 1 nM LT3, 500 μM IBMX, 100 ng/mL BMP7 (R&D, 354-BP) and 10 μM SB-431542. Medium was changed every other day.

#### White adipocyte differentiation

For WA differentiation, adipogenic MSCs were first cultured in B8 medium for 9 days, after which the medium was switch to W4 medium for another 21 days. The **W4 medium** made of DM was supplemented with 200 ng/mL IGF, 1 μM dexamethasone, 1 nM LT-3, and 10 μM SB-431542. Medium was changed every other day.

### EBF2-mScarlet/UCP1-tagGFP and TGFBR2-LOF line

The UCP1-tagGFP reporter H9-hESC line was previously created by Dr. Liang Zhang at University of Georgia. To knock in the IRES-mScarlet cassette to the 3’ UTR of EBF2 locus, the sgRNA target sequence was cloned into the BbsI site of px330 vector (Addgene, 42230). The donor plasmid for homology directed repair (HDR) was generated by cloning 0.66 kb homology sequence flanking IRES-mClover3-polyA tail sequence in a pUC57 vector backbone using synthesis service from BGI Genomics.

The PAM sites for the sgRNA were mutated on the donor plasmid. To transfect H9-hESCs, 9 μg vectors (px330: pRGS surrogate reporter plasmid: HDR donor plasmid = 2:3:1) were electroporated into 3 x 10^6^ cells using a Neon electroporation system (voltage = 1050 V, width = 30 ms, pulse = 2 cycles; Thermo, MPK10025). RevitaCell (1:200; Thermo, A2644501) was supplemented during electroporation to increase viability. Two days after electroporation, single cells were flow sorted based on the surrogate reporter signals, and seeded onto pMEF (Sigma, PMEF-NL-P1) coated 96-well plates. 10 μM ROCK inhibitor (Aladdin, Y412681) or 1x in-house CEPT^95^ was used during sorting to enhance cell viability. Single cell-derived clones were cultured in HAIF and selected by genotyping PCR.

For TGFBR2-LOF line, gRNAs targeting intronic sequences flanking exon5 of TGFBR2 were used together with a surrogate reporter vector. Electroporation, single cell sorting and genotyping procedures were the same as previously described. All oligos for sgRNA targets and genotyping primers are listed in the Key Resource Table.

### Immunostaining and imaging

#### Sample collection and preparation

Cells cultured on Geltrex-coated 4- or 8-well chamber slides (ibidi) were washed with DPBS (Service Bio, G4200), fixed in 4% paraformaldehyde (PFA; Aladdin, C104190) for 20 minutes at room temperature, and stored in DPBS with 1:500 ProClin 300 (Sigma Aldrich, 48917-U) at 4 °C for further staining.

Fixed somite spheroids were dehydrated in 30% sucrose (Sigma, S0389) overnight at 4 °C, then placed into 1 cm × 1 cm cryomolds (Tissue Tek, 4566), and embedded in O.C.T. compound (Tissue Tek, 4593). The snap-frozen samples were stored at −80 °C, until sectioned at 10 µm on a cryostat (Leica CM1950).

#### Immunofluorescent microscopy

Fixed samples were retrieved with pre-warmed (55 °C) 0.5% SDS/DPBS (Thermo Fisher, AM9820) for 15 minutes at room temperature, washed twice with DPBS, and permeabilized in 0.5% Triton X-100/DPBS (Sigma, T8787) for 30 minutes. After washing with 0.04% Tween 20/DPBS (Solarbio, T8220), non-specific antigens were blocked using Duolink Blocking Solution (Sigma, DUO82007) for 1 hour at room temperature. Primary antibodies (diluted in MAXbind Staining Medium; Active Motif, 15253) were incubated with samples overnight at 4 °C. On the next day, following three 5-minute washes with MaxWash Buffer (Active Motif, 15254), secondary antibodies (Thermo) and DAPI (TCI, D5888) in MAXbind were incubated with samples for 1 hour at room temperature. After 3 washes using DPBS, ProLong Gold Antifade (Thermo Fisher, P36934) mounting medium was applied to preserve fluorescent signals. Images were captured on an EVOS M5000 or Keyence All-in-One platform and processed with manufacturer software. Quantification of signal intensities were performed using ImageJ (https://imagej.net/). Antibodies used in this study are listed in the Key Resource Table.

### Immunohistochemical staining of human trunk embryoids (hTEM)

To dissect the distinct compartment layers of endotome and dermomyotome in human somtie trajectory, we re-created our previously developed human trunk embryoid version 3 (hTEM.v3).^18^ Briefly, day-6.5 hTEM.v3 samples were OCT-embedded and transversely sectioned at 10 µm on a Leica cryostat. Primary antibodies of anti-SOX2, anti-EBF2 and anti-PAX7 were used to label neural tube, endotome and dermomyotome, respectively. The immunostaining of sectioned slides and imaging procedure were as described in previous section. Antibodies used in this study are listed in the Key Resource Table.

### RNA extraction and real-time qPCR

RNA extraction was performed using the Total RNA Kit I (Omega, R6834) or Kit II (Omega, R6934), depending on the lipid composition of the cells. RNA quality and purity were assessed using a SYNERGY HTX multi-mode reader (BioTek). For reverse transcription, 1ug of total RNA was used in a 20ul reaction volume to synthesize cDNA templates using iScript cDNA Synthesis Kit (Bio-Rad, 1708841). Quantitative RT-qPCR was performed on a QuantStudio 7 Pro Real-Time PCR system (Applied Biosystem, A43182) using the Taqman Universal Probe Master Mix (Vazyme, QN113-01) with TaqMan probe sets (Thermo). The relative expression levels of mRNAs were normalized 18S rRNA using the ΔΔCt method. mRNA levels were plotted as mean ± SEM values. Taqman probes used in this study are listed in the Key Resource Table.

### Flow cytometry

Cells were dissociated into single cells using Accutase except hESC-derived adipogenic MSCs were dissociated using ACF enzymatic dissociation solution (StemCell Technologies, 05426). Then cells were washed DPBS and centrifuged at 200 x g for 4 minutes. For direct flow cytometry, cells were resuspended in fresh culture medium. For surface marker staining, freshly harvested cells were incubated with fluorescence-conjugated antibodies on ice for 30 minutes in dark. These stained cells were washed twice with ice-cold DPBS before analysis using a BD FACSAria Fusion (BD Biosciences). Flow data were analyzed using Flow Jo (v10). Flow antibodies used in this study are listed in the Key Resource Table.

### Lipid (Oil Red O) staining assay

To assess the lipid droplet formation potency upon treatments, BAs of endotome lineage were differentiated in TC surface 96-well plates (Corning, 3599) and fixed with 4% PFA at day 33 for Oil Red O staining, according to manufacture instructions (Sigma, MAK194). Briefly, fixed cells were washed with 60% isopropanol, then incubated with 100 ul of 3 mg/ml Oil Red O solution in 60% isopropanol for 15 minutes at room temperature. After incubation, stained cells were rinsed 2 to 5 times with distilled water, until the background was clear.

Brightfield images were captured per well using a BZ-X All-in-One platform (4× objective, stitch mode). The Oil Red O threshold for the region of interest was determined by visual inspection of clearly delineated lipid droplets. Stitched whole-well images were analyzed to calculate the percentage of Oil Red O-positive area from each well.

### CUT&Tag library preparation

Cells from day-6 somite and day-8 endotome cultures were dissociated into single-cell suspensions using Accutase. Then the cells were counted using a hemocytometer and processed according to manufacturer’s instruction of CUT&Tag assay kit (CST, #77552) with slight modifications. In brief, 100,000 live cells per experiment were suspended with kit-provided Concanavalin A beads in the binding buffer. Next, 2.5 ug primary antibody per sample was incubated with bead-coupled cells at 4°C overnight. The antibodies used were sheep polyclonal anti-EBF2 (R&D, AF7006) and rabbit polyclonal anti-H3K27ac (active motif, #39133). On the next day, 1 ul secondary antibody (donkey anti-sheep IgG was separately purchased from Sigma) per sample was added to the mixture and incubated at room temperature for 30 minutes. Next, 2 ul pAG-Tn5 was added to each sample and incubated at room temperature for 1 hour. Tagmentation was initiated by replacing the High Salt Digitonin Buffer with the Tagmentation Buffer containing MgCl_2_. After tagmentation and reverse crosslinking as instructed by the manufacture, tagmented DNA fragments were recovered and purified using DNA Clean & Concentrator-5 columns (Zymo, 77552S). We directly performed PCR amplification of the library from the eluted DNA in DNase/RNase-free water (Thermo, 10977023), as described for ChIPmentation protocol.^96^ The library PCR was set up by adding 25 ul NEBnext high-fidelity 2X PCR master mix (NEB, M0541S), 2.5 ul of each adapter primer, 15 ul eluted DNA and was amplified for 12 cycles (98 °C 10 s, 63 °C 30 s, 72 °C 30 s). The adapter primers used for each library was listed in the Key Resource Table. The amplified libraries were purified with 1.3 volumes of Mag-Bind TotalPure NGS beads (Omega, M1378). The libraries were quantified using Tapestation D1000 DNA analysis and sequenced with Nextseq 2000 (illumina) with the paired-end mode.

### CUT&Tag data processing

Raw single-indexed fastq reads were aligned to the human genome reference hg38 using STAR (v2.7.10a) with default settings. The resulting sorted BAM files were indexed with samtools (v1.9),^97^ and converted to bigwig format using bamCoverage function from deepTools (v3.5.1) with following options –binSize 50, --normalizeUsing RPGC, --ignoreForNormalization chrX chrY chrM, --ignoreDuplicates, --extendReads 150. The generated bigwig files were visualized with Integrative Genome Viewer (IGV) (v2.16.0).

To profile signals EBF2 and H3K27ac signals at putative enhancers in endotome cells, we first identified H3K27ac-enriched (foldenrichment > 5) regions in day-8 endotome using macs2 (v2.2.6)^98^ and annotate them to nearby genes with HOMER (v4.9.1). This gene list was then intersected with top1,000 differentially expressed genes from day-8 endotome scRNA-seq dataset to define the putative enhancers. Finally, we used bamCoverage and plotHeatmap functions from deepTools^99^ to visualize the differential signals of EBF2 and H3K27ac between day-6 somite and day-8 endotome cells.

### Single cell RNA sequencing

#### Preparation of single cell suspension

PAX3-mClover3/EBF2-mScarlet H9-hESC line was differentiated into somite, BAs of endotome, BAs of dermomyotome, vSMCs of endotome, and adipogenic MSC-derived beige adipocytes as described above. Cultures were harvested at indicated time points in Figures S5A, S5B and 6E.

For routine cultures, monolayered cells were dissociated from TC surface using Accutase at 37 °C for 5-10 minutes. For densely packed brown (endotome-derived BA at day24, dermomyotome-derived BAs at day 36 and 46) and beige adipocyte cultures, which were rich in extracellular matrix and lipids, cells were detached using 2.5 mg/ml collagenase IV (Thermo, 17104019) and 0.25% Trypsin-EDTA (Thermo, 25200056) at 37 °C for 10-15 minutes. The resulting brown and beige adipocyte suspensions were filtered through 100 µm strainers, centrifuged at 1,000 rpm for 4 minutes, and resuspended in DPBS at a density of 1 x 105 cells/ml in DPBS for single-cell capture using Singleron platform.

#### Single cell sequencing

scRNA-seq was performed on Singleron platform according to the manufacturer’s instructions. Briefly, single-cell suspensions were loaded into microfluidic chips using the Singleron Matrix^®^ Single-Cell Processing System (Singleron). scRNA-seq libraries were subsequently constructed following the manufacturer’s protocol for the GEXSCOPE^®^ Single-Cell RNA Library Kit (Singleron). Individual libraries were diluted to 4 nM, pooled, and sequenced on an Illumina HiSeq X instrument with the PE150 mode.

### Single cell sequencing data analysis

#### Raw data processing and quality control

scRNA-seq data generated in this study: For data generated using the Singleron platform, raw reads were processed with the SCOPE-tools pipeline (https://github.com/SingleronBio/SCOPE-tools) and aligned to the GRCh38 reference genome (Ensembl v99). Datasets were subsequently processed using the Seurat pipeline (v4.3.0.1) in R (v4.3.3). Mitochondrial and ribosomal UMI percentages were calculated with the PercentageFeatureSet function. Low-quality cells were filtered out based on the following criteria, fewer than 1,000 or more than 5,000 detected genes, mitochondrial transcript percentages outside the 3-15 % range, or total RNA counts greater than 15,000. Potential doublets and ambient RNA contamination were detected and labeled using the MarkDoublets and RemoveAmbientRNA functions from the scutilsR package (v0.1.1). The filtered expression matrix was normalized using NormalizeData, and the top 2,000 highly variable genes were identified using FindVariableFeatures for subsequent scaling. Cell cycle scores (S and G2/M phases) were calculated with CellCycleScoring and regressed out before principal component analysis (PCA) and UMAP embedding.

Public scRNA-seq data used in this study: hTEM.v3 dataset previously generated in our lab were downloaded from GSE314260. Mouse organogenesis datasets were obtained from the MouseGastrulationData package (v1.20.0). Cells representing human (days 5-7) and mouse (E7.25-E9.5) somite trajectories were extracted and converted to Seurat objects for downstream analysis. *in vitro* human dermomyotome-derived BAs and *in vivo* mouse interscapular BAT datasets were from GSE185623 for comparative analysis. *in vivo* human subcutaneous adipose tissue dataset was from GSE176171, *in vivo* human and mouse peri-aortic BAT datasets were from GSE164528. Cell type annotations were from their original metadata.

#### Batch effect correction and integration analysis

After quality filtering as described in the previous section, 72,927 cells were retained for BA development of endotome (day 6/8/12/16/20/24), 81,800 cells for BA development of dermomyotome (day 6/8/13/20/26/36/46), and 36,601 cells for endotome-derived vSMC (day 13/20). Prior to batch correction, cell cycle scores and mitochondrial/ribosomal percentages were regressed out using the ScaleData function to minimize cell-state-related variation. Integration for each pathway was performed using bbknnR^100^ (https://github.com/ycli1995/bbknnR) (v2.0.2) with method = bbknn and culture time (day) as the batch factor. Following integration, cells were clustered using the Louvain algorithm on the constructed bbknn graph. Differentially expressed genes between clusters were identified using FindMarkers with min.pct = 0.1 and logfc.threshold = 0.1.

#### Pseudotime analysis

The scaled data layer from the Seurat object was converted to .h5ad format using convertFormat function in sceasy (v0.0.7). Pseudotime analysis was performed with Palantir (v1.3.3)^101^ in a Python 3.9 environment using default parameters, with PAX3 positive somite cells pre-defined as the developmental starting point. Pseudotime trajectories representing divergent differentiation of endotome and dermomyotome were visualized using UMAP and force-directed graph (FDG) layouts, respectively. The inferred pseudotime values were subsequently added to the metadata of the corresponding objects for downstream analyses.

#### Gene ontology (GO) and gene set enrichment analysis (GSEA)

GO and GSEA were performed using clusterProfiler (v4.14.6, https://github.com/YuLab-SMU/clusterProfiler) in R. For GO analysis, cluster-specific genes were converted to Entrez Gene IDs using org.Hs.eg.db (v3.20.0). Biological process (BP) enrichment was conducted with the enrichGO function. Results were visualized using heatplot and barplot using Enrichplot (v1.26.6, https://github.com/YuLab-SMU/enrichplot) combined with ggplot2 (v3.5.1). For GSEA, genes were ranked by log₂(fold change), and the ranked list was analyzed using the GSEA function to identify significantly (-log_10_(p value) > 1) enriched pathways. GSEA results were visualized with the GseaVis package (v0.1.1).^102^

#### Mouse-to-human gene conversion and transcriptomic correlation analysis

Orthologous gene conversion from mouse to human was performed using the useMart and getLDS functions in biomaRt (v2.62.1). To compute the expression correlation in Figure 1H, cor function in R was used with method = Pearson. The significance of correlation was calculated with Student’s *t-test*, and the results were visualized using corrplot (v0.95, https://github.com/taiyun/corrplot). pheatmap (v1.0.12, https://github.com/raivokolde/pheatmap) was used with correlation method = pearson. The significance of correlation was calculated with Student’s *t-test*.

To compare the developmental similarity among in vitro and in vivo BAs in Figure 4J, cells in clusters of ‘BAs’, ‘UCP1-expressing cells’ and ‘beige adipocyte’ were extracted from mouse interscapular BAT (GSE185623), mouse peri-aortic BAT (GSE185623), human peri-aortic BAT (GSE185623), human subcutaneous adipose tissue (GSE176171), MSC-derived beige adipocytes (this study), and BAs of endotome and dermomyotome (this study), were integrated into a single Seurat object for transcriptomic correlation analysis. Data were regressed for cell cycle scores and percentages of mitochondrial and ribosomal RNAs, using ScaleData. Batch effects were corrected using bbknnR, with batch factors defined as cell type and sequencing platform. A Euclidean distance matrix was calculated, and complete-linkage hierarchical clustering was applied to group samples by similarity. The resulting correlation matrix was visualized using the pheatmap package (v1.0.12).

### Seahorse analysis

#### Extracellular flux assay

Seahorse assay was performed as previously.^3^ Briefly, 10,000 cells from day-12 endotome-BA lineage and from day-35 adipogenic MSCs were seeded onto geltrex coated Seahorse XFe96 cell culture microplates (Agilent, 103793-100). For parallel brown and white adipogenesis, cells were culture in BA and W4 medium, respectively, for 21 days (see previous sections of BA and WA differentiation). Twenty-four hours prior to the flux assay, cells were given freshly prepared flux assay medium. The flux assay medium was composed of XF base medium (Aglient, 102353-100), 25 mM glucose (Sigma, G5767), 1x sodium pyruvate (Thermo, 11360039) and 1x GlutaMAX. 10 µM Forskolin (Tocris, 1099) or DMSO vehicle were added to the XF assay medium to measure changes in respiration rates. Extracellular acidification rates (ECAR) and oxygen consumption rates (OCR) were measured using the Seahorse XF Cell Mito Stress Kit (Agilent, 103015-100) on a Seahorse XFe96 Analyzer (Agilent), following the manufacturer’s instructions. Oligomycin (2 µM), FCCP (2 µM), and rotenone/antimycin A (5 µM) were injected at the indicated time points. ECARs and OCRs were measured from 8 technical replicates.

To normalize the respiration capacity for differences in cell number between BA and WA cultures, cells were lysed immediately after flux analysis, and total genomic DNA was quantified from each well using the EasyPure Genomic DNA kit (TransGen Biotech, EE101-01) and a SYNERGY HTX multi-mode reader. Raw flux data were processed with WAVE software (Agilent), and the resulting OCR and ECAR values were normalized to corresponding DNA content per well.

#### Fatty acid oxidization assay

To assess the capacity for extracellular fatty acid oxidation, both BAs and WAs were incubated in substrate-limited medium for 24 hours prior to the flux assay. This medium consisted of glucose-free DMEM (Thermo, A1443001) supplemented with 1× sodium pyruvate, 0.5 mM glucose, 1× GlutaMAX, 0.5 mM carnitine (MCE, HY-B0399) and 2% FBS. Meanwhile, 4 µM Etomoxir (TargetMol, T4535) or DMSO vehicle was added.

Immediately before the flux assay, the medium was replaced with XF base medium containing 5.5 mM glucose, 0.5 mM carnitine and either 150 µM palmitate-BSA (Agilent, 102720) or BSA alone. ECAR and OCR measurements. Raw data normalization to DNA content, were performed as described in the previous section.

### Quantification and statistical analysis

Standard statistical analyses were performed using GraphPad Prism (v10). All data are presented as mean ± standard error of the mean (SEM). Statistical significance between two groups was determined using an unpaired two-tailed Student’s *t*-*test*, and comparisons among three or more groups were performed using one-way ANOVA followed by Bonferroni’s post-hoc test. Differences were considered statistically significant at *p* < 0.05 (*), *p* < 0.01 (**), *p* < 0.001 (***), and *p* < 0.0001 (****).

